# Topological analysis of single-cell data reveals shared glial landscape of macular degeneration and neurodegenerative diseases

**DOI:** 10.1101/2021.01.19.427286

**Authors:** Manik Kuchroo, Marcello DiStasio, Eda Calapkulu, Maryam Ige, Le Zhang, Amar H. Sheth, Madhvi Menon, Yu Xing, Scott Gigante, Jessie Huang, Rahul M. Dhodapkar, Bastian Rieck, Guy Wolf, Smita Krishnaswamy, Brian P. Hafler

## Abstract

1

A novel topological machine learning approach applied to single-nucleus RNA sequencing from human retinas with age-related macular degeneration identifies interacting disease phase-specific glial activation states shared with Alzheimer’s disease and multiple sclerosis.

**Abstract:** Neurodegeneration occurs in a wide range of diseases, including age-related macular degeneration (AMD), Alzheimer’s disease (AD), and multiple sclerosis (MS), each with distinct inciting events. To determine whether glial transcriptional states are shared across phases of degeneration, we sequenced 50,498 nuclei from the retinas of seven AMD patients and six healthy controls, generating the first single-cell transcriptomic atlas of AMD. We identified groupings of cells implicated in disease pathogenesis by applying a novel topologically-inspired machine learning approach called ‘diffusion condensation.’ By calculating diffusion homology features and performing persistence analysis, diffusion condensation identified activated glial states enriched in the early phases of AMD, AD, and MS as well as an AMD-specific proangiogenic astrocyte state promoting pathogenic neovascularization in advanced AMD. Finally, by mapping the expression of disease-associated genes to glial states, we identified key signaling interactions creating hypotheses for therapeutic intervention. Our topological analysis identified an integrated disease-phase specific glial landscape that is shared across neurodegenerative conditions affecting the central nervous system.

## 3 Introduction

Age-related macular degeneration (AMD) is a progressive neurodegenerative disease of the retina that affects 170 million individuals worldwide and is a leading cause of blindness in the elderly [*1*]. Similar to Alzheimer’s disease (AD) and progressive multiple sclerosis (MS), AMD pathology can be categorized into distinct stages. Initially, in early, dry AMD, focal deposits of extracellular lipid-rich debris known as drusen accumulate below the retinal pigment epithelium leading to activation of glia [*2*]. In advanced, neovascular AMD, the development of angiogenesis and fibrosis causes photoreceptor loss and a progressive decline in central visual acuity [*3*]. In MS and AD, glial dysregulation is associated with neuronal damage and progressive neurologic impairment [*4, 5*]. While application of single-cell transcriptomics has given insight into the cell states enriched in AD and MS [*5, 6, 7, 8*], a single-cell analysis of retinal tissue across phases of AMD disease progression has not yet been performed and integrated into our understanding of the common mechanisms of neurodegeneration. To identify cell types and states enriched across AMD phases, we performed massively parallel microfluidics-based single nucleus RNA-sequencing (snRNA-seq) to create the first single-cell atlas of AMD pathology.

While recent application of snRNA-seq to AD and MS has identified disease-associated microglial and astrocytic states [*5, 6, 8, 7*], identifying related states in the retina has remained a challenge [*9, 10*]. Existing clustering approaches present only one or a few levels of the cellular hierarchy, often missing information such as rare cell types or salient populations affected by disease status [*11, 12, 13*]. Furthermore, current approaches enforce global granularity constraints which can hamper disease comparisons across tissues, since the optimal level for one tissue may obscure similarities in cellular substructure in another tissue. To address these concerns, we combined our single cell dataset with a novel topologically-inspired machine learning tool called diffusion condensation, which sweeps through different granularities of data to identify natural groupings of cells at each level. While clustering tools partition data into one or a few levels of granularity, diffusion condensation constructs a full hierarchy of cellular states, or a diffusion homology, which can be evaluated with persistence analysis to identify cell types, subtypes and disease-phase associated cellular states. Furthermore, by comparing key cellular groupings across tissues, diffusion condensation can help identify related cell states apparent at disparate granularities.

With our combined approach, we integrated diffusion condensation results from AMD, AD and MS snRNA-seq datasets to identify shared activation states in glia in the early phase of all three neurodegenerative diseases. Using interaction analysis, we identified key signaling interactions between activated glial states as well as a proangiogenic astrocyte state enriched in late-stage AMD, a likely driver of late-stage neovascularization and photoreceptor loss [*3*]. Our study defines the transcriptional landscape in AMD and reveals key phase-specific glial states shared across multiple neurodegenerative diseases affecting the brain and retina.

## 4 Results

### Massively parallel single-nucleus sequencing of AMD and control retinas

The retina and the brain are complex central nervous system (CNS) tissues with many hierarchies of cell types and subtypes which disease status can differentially affect (Fig. 1A). As a component of the CNS, the retina shares features with the brain at the level of cell biology and degenerative pathology (Fig. 1B). Similar to AMD, MS and AD have defined disease phases, each with an early or acute active, and a late or chronic inactive disease stage [*14,15,16*]. To identify cellular states enriched in AMD, we performed massively parallel microfluidics-based single-nucleus RNA-sequencing (snRNA-seq) on 50,498 isolated retinal nuclei from the maculas of six postmortem healthy control retinas and seven with varying stages of AMD pathology, creating the first single-cell dataset of AMD pathology.

**Figure 1:**
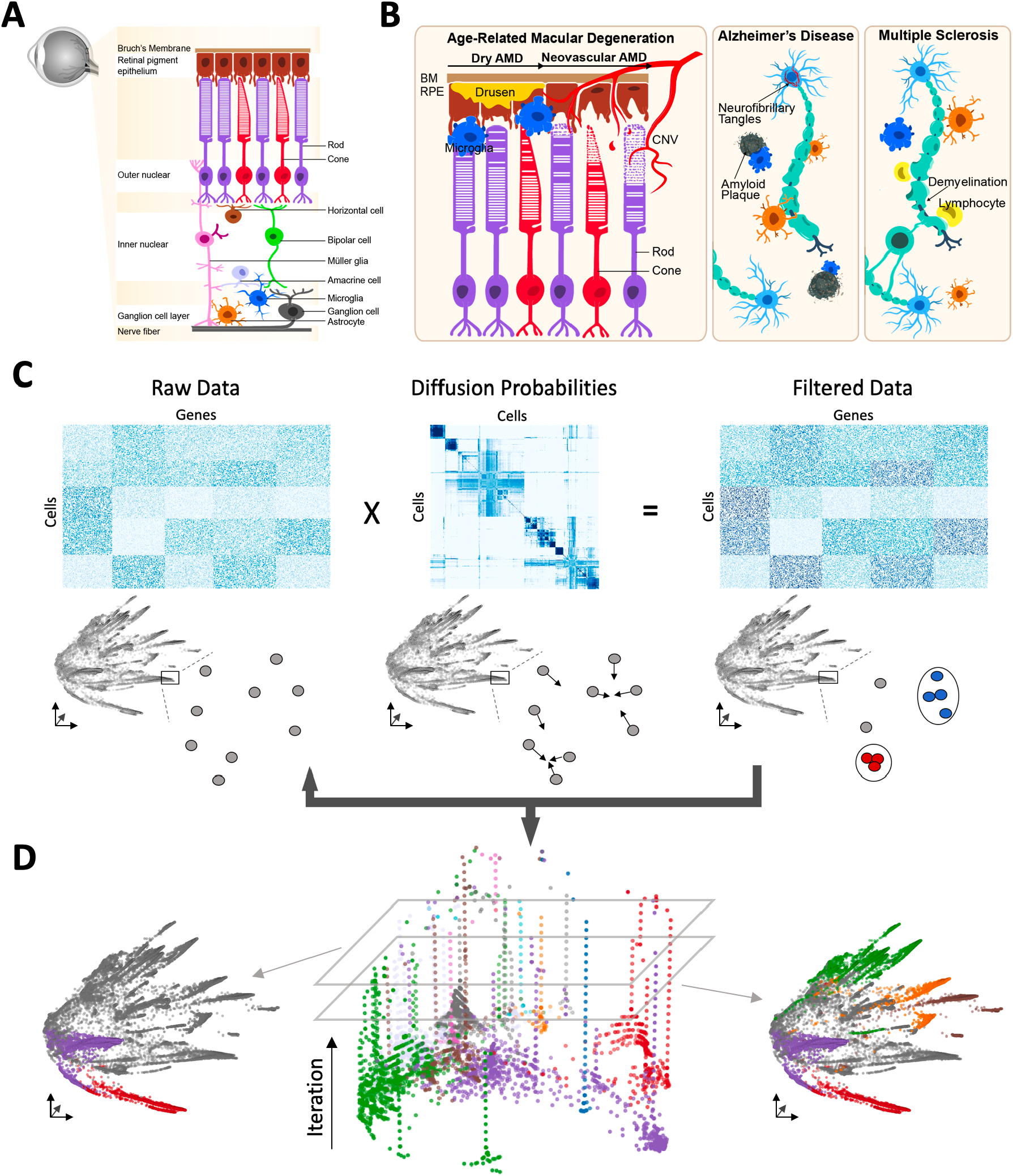
Overview of neurodegenerative disease processes and the topological diffusion condensation approach. **(A)** Sketch of retina cross section showing layers and major cell types. **(B)** Illustration of the role of innate immune cells in neurodegenerative disease pathogenesis. In the dry stage of AMD there is accumulation of extracellular drusen debris below the retinal pigment epithelium and activation of innate immune cells. **(C)** The diffusion condensation process for building a topological understanding of the single cell dataset. Assigning cells **[left]** to topological features across multiple granularities involves repeated iteration of three successive steps. The first step computes a diffusion operator **[center]** from the input data. Next, the diffusion operator is applied to the data set to produce a filtered output. Finally, cells that fall below a threshold *ζ* are merged into a single group or topological feature at a particular granularity **[right]**. These steps are repeated iteratively till all cells have merged into a single feature. **(D)** Visualization of diffusion condensation’s diffusion homology calculation. Evaluating persistent topological features, or groupings of similar cells, in the diffusion homology identifies clusters of cells with similar transcriptional profiles across granularities.

To identify disease-associated transcriptional states, we applied the community-detection based multigranular clustering tool Louvain to the AMD single-cell transcriptomic dataset [*17*]. Louvain revealed 22 populations at coarse granularity, and 40 populations at fine granularity (Supplementary Fig. 1A-B). Across both resolutions, however, rare innate immune cell types such as microglia, astrocytes and Müller glia, were not identified, with genes specific for these cells types not localizing to any one cluster. While these cell type specific signatures were not isolated by Louvain clusters across granularities, they did localize in the PHATE embedding, revealing that they are geometrically isolated on the cellular manifold [*18*] (Supplementary Fig. 1C).

### Diffusion condensation identifies biologically meaningful groupings of cells across multiple levels of granularity

The difference between diseased and normal tissue can appear as systemic differences in large populations of neurons or as subtle shifts in rare cell types of the innate immune compartment. One purpose of single cell data is to identify meaningful subpopulations and transcriptional states in an unbiased fashion from diseased and normal tissue. However, while biological data has structure at many different levels of granularity, most clustering methods offer one or just a few levels of granularity. These few levels of granularity can create inaccurate identifications of disease-associated cellular states. To address this, we develop a method that combines the principles of data manifold geometry with computational topology to create a better understanding of cellular states across granularities.

Geometric analysis on a cellular manifold is based on a notion of *distances* between different cells. Previous work in data diffusion has shown that this approach can be useful for both visualization and clustering [*19, 18*], revealing the overall structure of the dataset. Topological analysis, on the other hand, is based on how these cells relate to one another in their local geometry as dictated by structure at various granularities. Topological data analysis, meanwhile, constitutes a hybrid framework between a purely geometrical view and a purely topological one [*20*]. Topological data analysis identifies related cells across different granularities (geometric scales) by first computing a pairwise distance matrix **D**, before identifying all cell pairs whose distance falls below a distance threshold *δ*. A pair of cells that fall below this threshold are deemed to be part of the same ‘connected component’ or ‘topological feature.’ The value of *δ* is akin to a granularity level at which we view the data: as *δ* increases, more cell pairs will be connected, quickly creating more connected components at coarser granularities. Persistent homology is a framework that tracks connected components that persist across a range of granularities. All cells belonging to such a component are members of the same cluster across multiple scales. Inspired by both persistent homology and the geometric approach to identify cell populations, we further developed diffusion condensation, a dynamic process in which data points slowly and iteratively condense towards local centers of gravity to create a topological understanding of what we refer to as diffusion homology [*10*].

Diffusion condensation uses the geometry of data to extract topological features from single cell data (Fig. 1C, D). While single cells are measured in high dimensional gene space, cellular states can be modeled in a substantially lower dimensional manifold which captures the geometric variability in the dataset. Data diffusion works to visualize this single cell manifold by computing *diffusion probabilities* between related cells [*19, 18*]. Previous applications of these *diffusion probabilities* to single cell data have revealed that data diffusion can also be used to filter cells, effectively moving them towards the center of a local neighborhood of related cells [*21*].

Diffusion condensation builds upon this previous work in topological data analysis, diffusion geometry and diffusion filters by applying these *diffusion probabilities* iteratively to the single cell dataset, effectively creating a topological understanding of the dataset based on its geometry. More precisely, the diffusion condensation process starts by computing a distance matrix **D** between all cells, before converting this to an affinity matrix **K** by using a fixed bandwidth Gaussian kernel function [*10*]. Next, **K** is row normalized to obtain *diffusion probabilities* between cells. These diffusion probabilities are applied to the input data to replace the value of a point with the weighted average of its diffusion neighbors. This process removes high frequency variation in terms of the graph spectrum of the cell-cell graph, effectively causing points to *condense*, or move closer together on the cellular manifold. In the third step, diffusion condensation merges points together that have condensed within a preset distance threshold *ζ*, effectively connecting two previously independent components or cells. This process is then repeated until all cells have collapsed to a single connected component. At each iteration, diffusion probabilities are calculated on the output of the previous iteration and used as a filter to produce an increasingly coarse-grained data geometry for the next iteration (Fig. 1C). This deep cascade of filters effectively builds a topology by connecting data points together in a natural manner rather than forcing merges as done by agglomerative clustering techniques (see Methods).

While the original description of diffusion condensation [*10*] only invoked diffusion geometry, here we recognize and extend its connections to topology. When applied to single cell data, we can analyze the resulting *diffusion homology*, a novel description inspired by persistent homology, which helps understand the evolution of topological features during the condensation process. The underlying idea is to summarize the merging of points during the condensation process and assigning each connected component a topological ‘prominence’ value known as *persistence*. Our analysis serves to identify topological features with a high persistence, focusing on 0D features also known as connected components. Highly persistent connected components are effectively groups of cells that are similar in their transcriptional profile. Through this analysis, we can identify major cell types, subtypes, and activation states across granularities. While we can visualize the overall diffusion homology (Fig. 1D), topological features and their associated persistence values are best visualized using a ‘persistence barcode.’ This is a visualization [*22*] consisting of horizontal bars of different lengths; each bar corresponds to one topological feature—a subgroup of cells in our case—while the length of each bar depicts the persistence of that feature, directly indicating to what extent the feature is prominent. The barcode serves as a summary of the condensation process but also makes it possible to easily select relevant scales or granularities at which to analyze the dataset.

While analysis of diffusion homology will identify populations of related cells in an unbiased manner, it will not identify topological features, and their underlying cellular populations, that are enriched in a disease condition of interest. In our analysis, cells from different diseases states, such as healthy, dry AMD and neovascular AMD, were included. To take this information into consideration, we use MELD to identify persistent topological features that are enriched or depleted in different disease phases [*23*]. MELD is a manifold-geometry based method of computing a likelihood score for each cell state, indicating whether it is more likely to be seen in the normal or disease sample. MELD works by creates a joint graph of both the normal and disease samples being compared, and returns a relative likelihood that quantifies how likely each cell in the population is of occurring in the disease versus normal condition. Combining this cell-level score with information from our topological analysis, we can identify granularities in the diffusion homology that optimally isolates cellular states enriched in differing disease conditions. A complete mathematical description and discussion of the novelty of the diffusion condensation algorithm and diffusion homology analysis can be found in the methods section, along with in-depth comparisons to other approaches applied to single cell data (see Methods).

### Diffusion condensation identifies all abundant and rare cell types in the retina

We applied diffusion condensation to the AMD snRNA-seq dataset to identify the major cell types present in the control and AMD samples. By analyzing the constructed diffusion homology, we identified persistent features in the barcode visualization, which corresponded to interconnected cellular populations (Fig. 2A). We categorized each of the unbiased populations based on the expression of previously established cell type specific marker genes [*24*] (Fig. 2B-C) (see Methods). Using this approach, we identified all neuronal cell types, including retinal ganglion cells, horizontal cells, bipolar cells, rod photoreceptors, cone photoreceptors, and amacrine cells, as well as all rare non-neuronal cell types, including microglia, astrocytes, Müller glia, and vascular cells.

**Figure 2:**
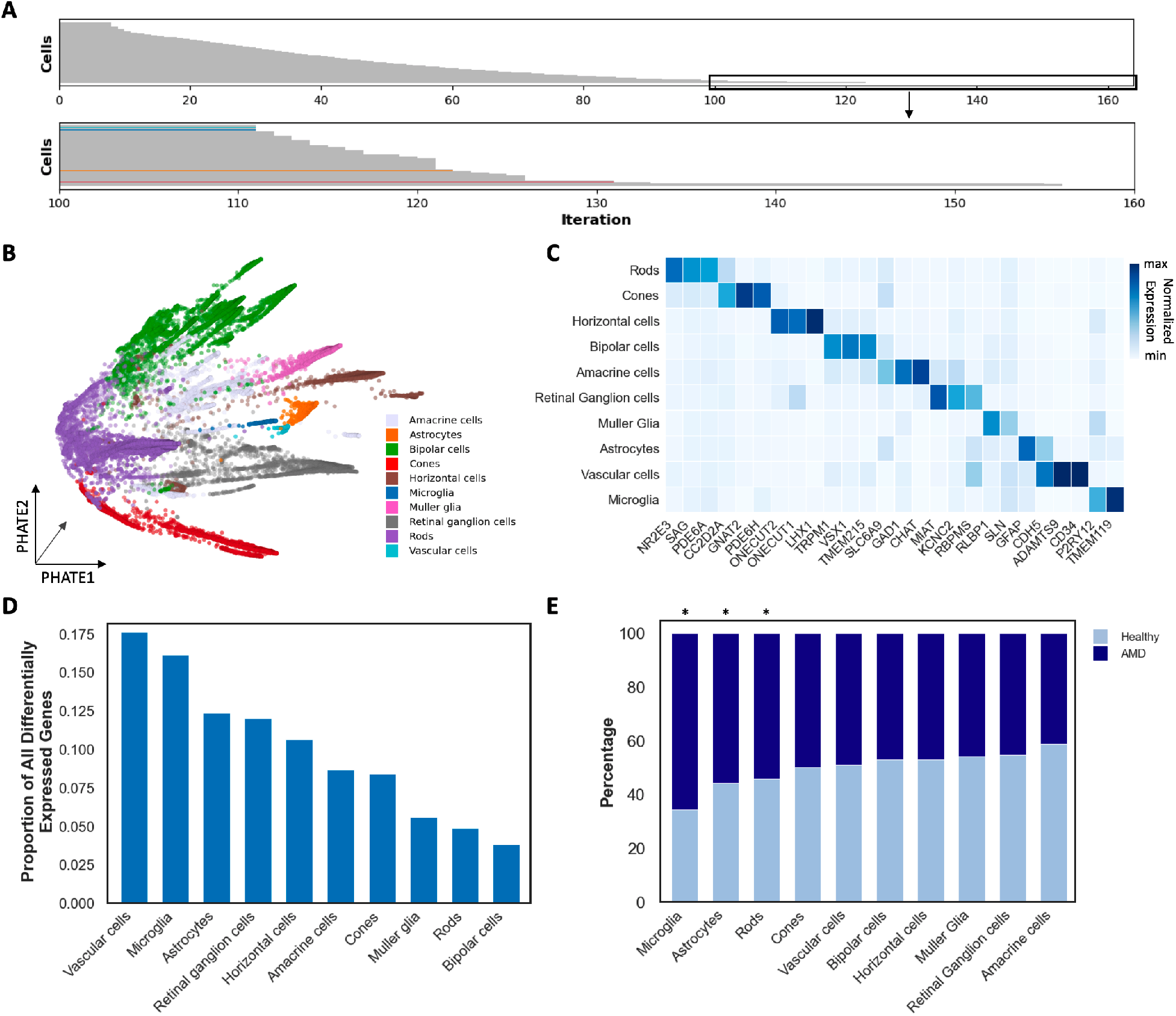
Diffusion condensation analysis of snRNAseq data from AMD tissue. **(A)** Visualization of persistence barcodes by applying diffusion condensation to 50,498 nuclei isolated from AMD and control samples. Barcode colors highlight persistent topological features that correspond with known cell types. **(B)** PHATE visualization of 50,498 nuclei isolated from AMD and healthy retinas. All major retinal cell types were identified by performing persistence analysis on the diffusion condensation computed diffusion homology [*18*]. **(C)** Diffusion condensation identified cell types, as shown by the average normalized expression of known cell type specific marker genes. **(D)** Differential expression analysis revealed that vascular cells, microglia and astrocytes have the most dysregulated genes in AMD samples over controls. **(E)** Bar chart indicates the contribution of cell types in each cluster from normalized control and AMD samples. Microglia and astrocytes are the most enriched cell types in AMD using cross-condition abundance analysis (p < 0.01, single-sided binomial test).

To demonstrate the ability of diffusion condensation to identify meaningful populations of cells across granularities, we further explored subtypes of bipolar cells. With 12 subtypes identified in the primate and human retina, bipolar cells define a diverse set of interneurons that transmits signals from rod and cone photoreceptors to retinal ganglion cells [*25, 26, 27*]. While a persistent cluster of bipolar cells was marked by the pan-bipolar cell maker *VSX1*, the first two major subtypes of bipolar cells, ON-center and OFF-center, were marked by *GRM6* and *GRIK1*, corresponding to the cells that depolarize and hyperpolarize, respectively, in response to light in their receptive field centers. Diffusion homology analysis revealed that the two most persistent topological features (connected components) within the bipolar cell feature, were the ON and OFF bipolar subtypes (Supplementary Fig. 3A). By analyzing the persistent topological features within the ON-center and OFF-center bipolar cells, we identified all 12 major subtypes of cells based on the expression of cell subtype specific marker genes (Supplementary Fig. 3B-D).

**Figure 3:**
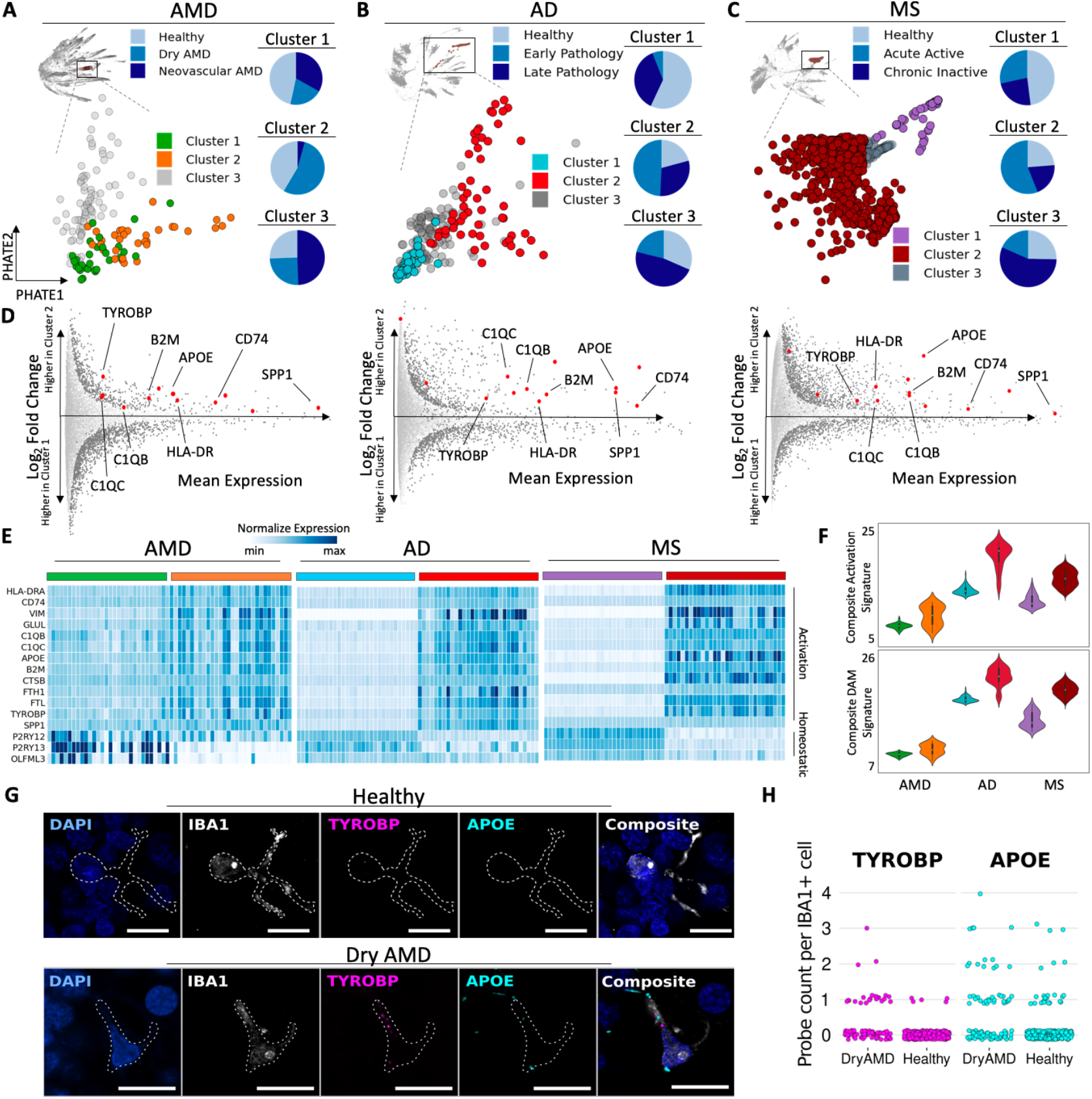
Analysis of microglia reveals a shared activation signature enriched in the early phase of three different neurodegenerative diseases. **(A)** With MELD, diffusion condensation isolated three populations of microglia in AMD and healthy retinas, each enriched for a different disease phase. Cluster 1: enriched for cells from healthy retinas; cluster 2: cells from dry AMD; cluster 3: cells from neovascular AMD. **(B)** As in panel A, three subsets of microglia are found in AD with diffusion condensation, each enriched for cells from a different disease condition. Cluster 1: enriched for cells from healthy brain tissue; cluster 2: cells from early disease state; cluster 3: cells from late disease state. **(C)** As in panel A, three subsets of microglia are found in MS with diffusion condensation, each enriched for cells from a different disease condition. Cluster 1: enriched for cells from healthy brain tissue; cluster 2: cells from an acute active lesion; cluster 3: cells from a chronic inactive lesion. **(D)** Differential expression analysis between cluster 1 (healthy-enriched) and cluster 2 (early or acute active disease-enriched) across all three degenerative diseases reveals a shared activation pattern in early disease. This signature includes *TYROBP*, *B2M*, *APOE*, *CD74*, *SPP1*, *HLA-DR*, *C1QB*, *C1QC*, *HLRA-DRA*, *VIM*, *GLUL*, *CTSB*, *FTH1*, *FTL*. **(E)** Heatmap demonstrating differences in expression of the neurodegenerative shared activation pattern and a homeostatic signature between cluster 1 and cluster 2 across all three degenerative diseases. **(F, upper)** Composite microglial activation signature for the neurodegenerative shared activation pattern in cluster 1 and cluster 2 across all three degenerative diseases. **(F, lower)** Disease associated microglia (DAM) signature [*7*] for cluster 1 and cluster 2 across all three degenerative diseases. **(G)** Micrographs of combined in-situ RNA hybridization and IBA1 immunofluorescence demonstrating elevated expression of key components of the neurodegenerative shared activation pattern (*TYROBP* and *APOE*) in IBA1-positive cells, a marker of microglia, from retinas with dry AMD as compared to controls. All scale bars = 10μm. **(H)** Quantification of expression of *TYROBP* and *APOE* in dry AMD retinas compared to controls.

To identify cell types implicated in AMD pathogenesis in an unbiased manner, we applied differential expression analysis to the diffusion condensation identified cell types. By comparing the cells from retinas with AMD to the cells from control retinas, we computed gene expression differences with Earth Mover’s Distance (EMD) within each cell type [*21*]. After setting a global significance cut off, we determined the number of differentially expressed genes by cell type. This analysis revealed that non-neuronal cell types in the retina, including vascular cells, microglia, and astrocytes, had the greatest number of differentially expressed genes between AMD and control samples (Fig. 2D). Furthermore, we performed abundance analysis to identify if certain cell types were significantly more enriched in AMD. This analysis revealed a significant increase in the normalized proportion of microglia and astrocyte nuclei from donors with AMD compared to control samples (Fig. 2E). These findings suggest that non-neuronal cell types including microglia and astrocytes are important cell types in AMD pathogenesis.

### Microglial activation signature identified in dry AMD is shared across the early phase of multiple neurodegenerative diseases

In order to identify microglial subpopulations enriched in specific disease phases of AMD, we identified topological features that isolated high MELD likelihood scores computed for healthy, dry and neovascular AMD conditions (Supplementary Fig. 4A) (see Methods). With this approach we identified three topological features or clusters, each enriched for a different condition: cluster 1 of microglia was enriched for cells from healthy controls, cluster 2 of microglia was enriched for cells from patients with dry AMD and cluster 3 of microglia was enriched for cells from patients with neovascular AMD (Fig. 3A). To identify signatures of AMD present in microglia during the early stage of disease pathogenesis, a phase in which microglia have been previously implicated [*2*], we performed EMD-based differential expression analysis between healthy-enriched cluster 1 and dry AMD-enriched cluster 2. Analyzing the top 100 most differentially expressed genes between these subpopulations, a clear activation signature appeared enriched in cluster 2, including genes known to play a role in neurodegeneration *APOE*, *TYROBP* and *SPP1* [*28*].

**Figure 4:**
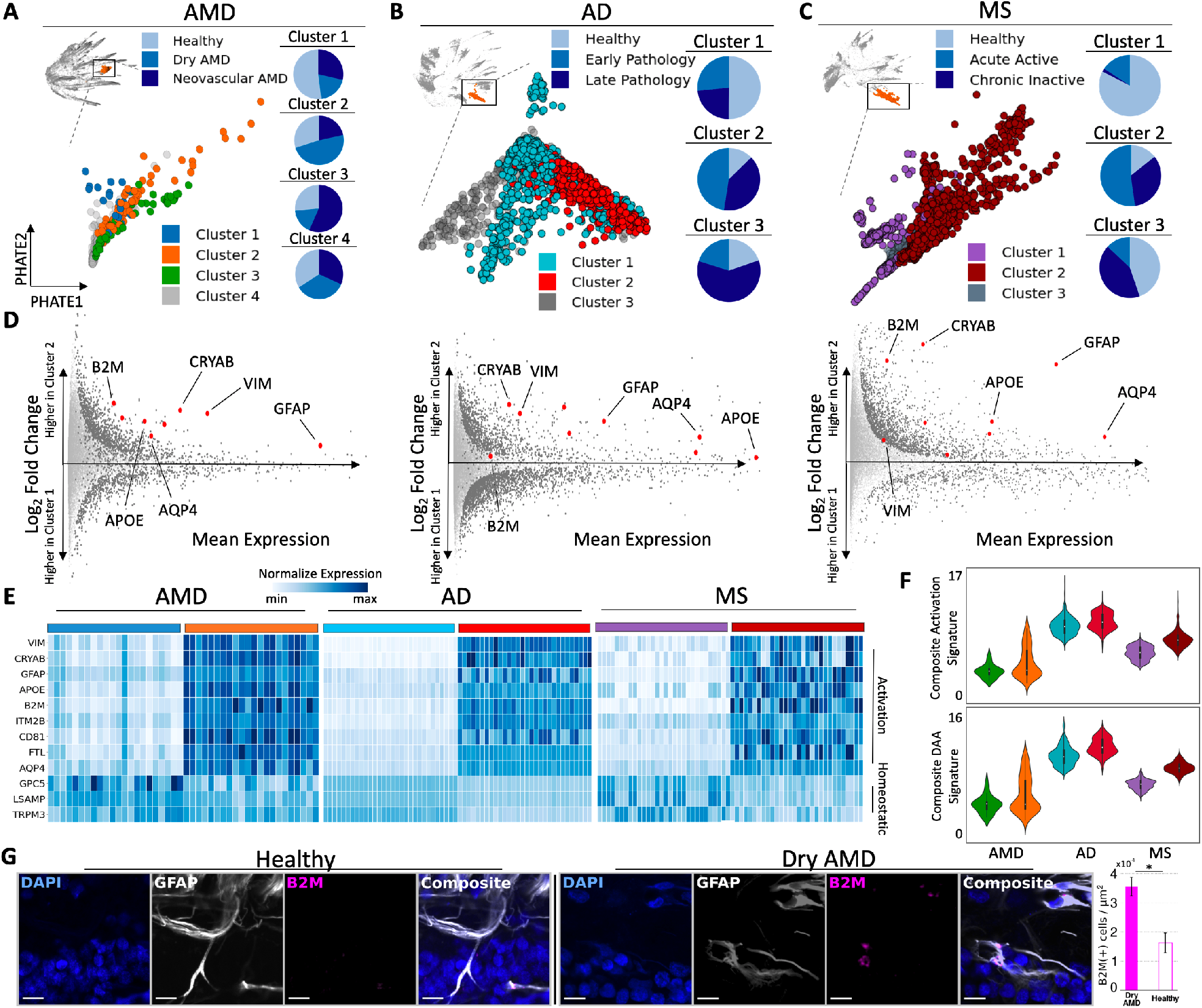
Analysis of astrocytes reveals a shared activation signature enriched in the early phase of three different neurodegenerative diseases. **(A)** With MELD, diffusion condensation isolated four populations of astrocytes in AMD and healthy retinas, each enriched for a different disease phase. Cluster 1: enriched for cells from healthy retinas; cluster 2: cells from dry AMD; cluster 3: cells from neovascular AMD; cluster 4: cells from all conditions equally. **(B)** As in panel A, three subsets of astrocytes are found in AD with diffusion condensation, each enriched for cells from a different disease condition. Cluster 1: enriched for cells from healthy brain tissue; cluster 2: cells from early disease state; cluster 3: cells from late disease state. As in panel A, three subsets of astrocytes are found in MS with diffusion condensation, each enriched for cells from a different disease condition. Cluster 1: enriched for cells from healthy brain tissue; cluster 2: cells from an acute active lesion; cluster 3: cells from a chronic inactive lesion. **(D)** Differential expression analysis between cluster 1 (healthy-enriched) and cluster 2 (early disease-enriched) across all three degenerative iseases reveals a shared activation pattern in early disease. This signature includes *B2M*, *CRYAB*, *VIM*, *GFAP*, *AQP4*, *APOE*, *ITM2B*, *CD81*, *FTL*. **(E)** Heatmap demonstrating differences in expression of the neurodegenerative shared activation pattern and a homeostatic signature between cluster 1 and cluster 2 across all three degenerative diseases. **(F, upper)** Composite astrocyte activation signature for the neurodegenerative shared activation pattern in cluster 1 and cluster 2 across all three degenerative diseases. **(F, lower)** Disease associated astrocyte (DAA) signature [*8*] for clusters 1 and cluster 2 across all three degenerative diseases. **(G)** Micrographs of combined in-situ RNA hybridization and GFAP immunofluorescence showing more abundant *B2M* expression in astrocyte-rich retinal layers from dry AMD retina when compared to healthy control. All scale bars = 10μm. Bar plot shows density of *B2M* transcripts in inner plexiform layer, retinal ganglion cell layer, and nerve fiber layer in retina samples affected by dry AMD and control.

Due to the similarity between this activation state with previously defined disease-associated microglia enriched in neurodegeneration [*7, 29*], we performed a comprehensive analysis of microglial states across other neurodegenerative diseases. We applied diffusion condensation to snRNA-seq data from AD [*4*] and MS [*30*] and analyzed each datasets diffusion homology features to identify all major cell types based on the expression of cell type specific marker genes (Supplementary Fig. 5A-D). Similar to AMD, abundance analysis revealed that microglia were significantly enriched in AD and MS when compared to healthy brain tissue (Supplementary Fig. 5E-F). To identify disease-phase specific transcriptomic states, we applied MELD to microglia in each dataset and identified three clusters of microglia in each disease: cluster 1, enriched for cells from healthy brain tissue; cluster 2, enriched for cells from early stage AD tissue or acute active MS lesions; and cluster 3, enriched for cells from late stage AD tissue or chronic inactive MS lesions (Fig. 3B,C). By comparing healthy enriched cluster 1 with early disease enriched cluster 2, a similar activation profile appeared in all three diseases (Fig. 3D). Integrating differential expression analysis results from all three diseases revealed a common signature of activation (*CD74*, *SPP1*, *VIM*, *FTL*, *B2M*), lipid and lysosomal phagocytosis (*APOE*, *TYROBP*, *CTSB*), and complement (*C1QB* and *C1QC*) (Fig. 3D). Furthermore, a common reduction was seen in the expression of homeostatic genes (*P2RY12*, *P2RY13*, and *OLFML3*) (Fig. 3E). We built a composite activation signature and disease-associated microglia signature [*7*] and mapped that onto our clusters, identifying that our early disease enriched clusters also displayed higher expression of the disease-associated microglia signature (Fig. 3F). This persistent topological feature comparison as performed by diffusion condensation significantly increased our ability to identify differentially expressed genes and develop a common signature (Supplementary Fig. 6A) (see Methods).

**Figure 5:**
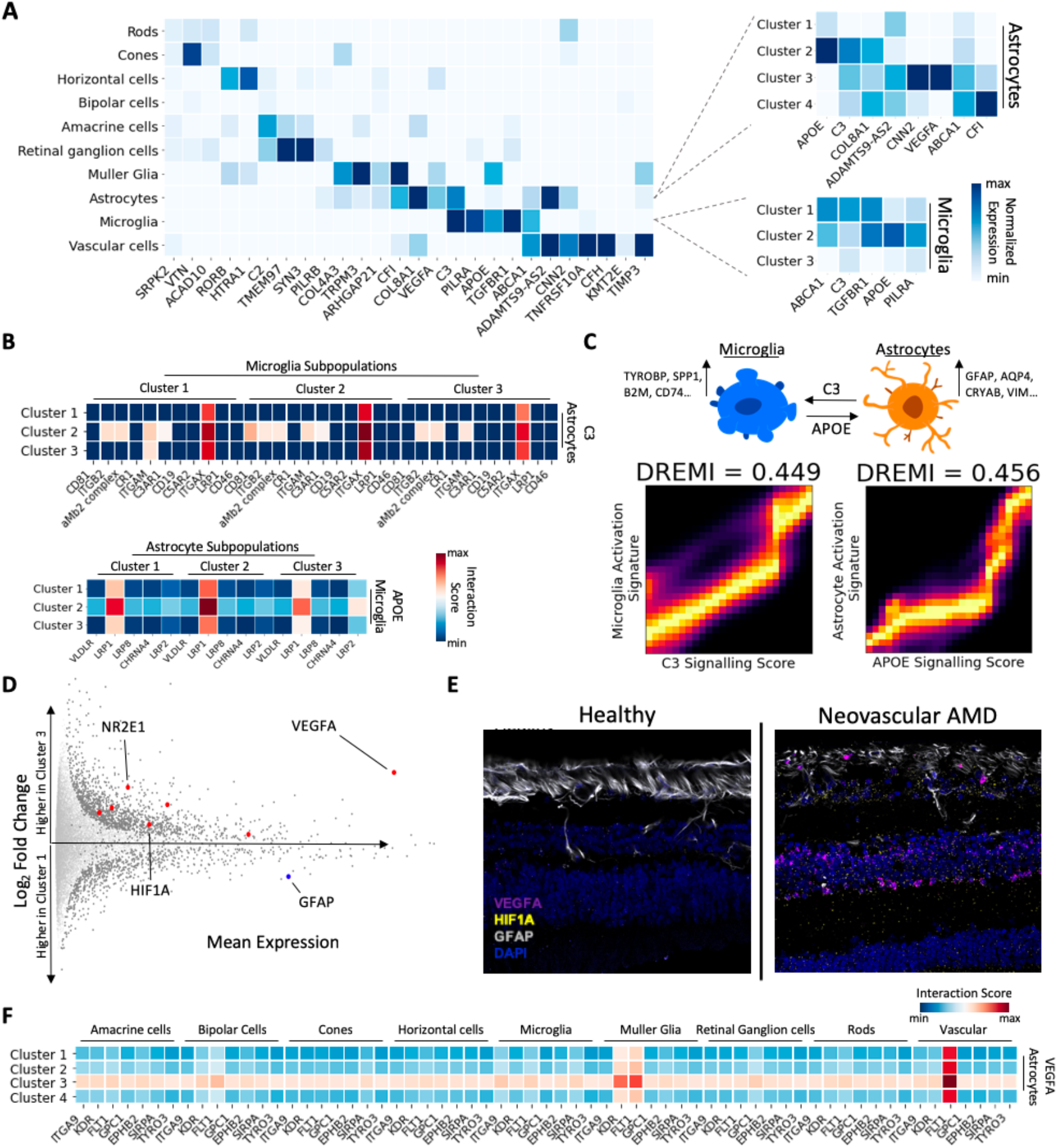
Mapping disease associated genes to diffusion condensation populations across granularities identifies potential signaling interactions. **(A)** Mapping AMD GWAS-associated genes to diffusion condensation identified cellular populations across granularities. **(B)** CellPhoneDB ligand-receptor interaction map between AMD-associated gene C3 expressed by the astrocyte subclusters and C3 receptor complex elements expressed by the microglia subclusters **[upper]**. CellPhoneDB ligand-receptor interaction map between the AMD-associated gene APOE expressed by the microglia subclusters and the APOE receptors expressed by the astrocyte subclusters **[lower]**. **(C)** DREVI visualizations of C3 and APOE signaling scores versus the composite early activation signatures (see Figs. 3 and 4) for microglia (left) and astrocytes (right). **(D)** Differential expression analysis between retinal data healthy-enriched cluster 1 and neovascular AMD-enriched cluster 3 reveals a pro-angiogenic signature present in advanced neovascular AMD (upregulated *VEGFA*, *NR2E1*, and *HIF1A* and decreased *GFAP*). **(E)** A marked increase in *HIF1A* and *VEGFA* expression is detected throughout retinal tissue in neovascular AMD disease state compared with healthy control visualized by in situ RNA hybridization, despite modest gliotic change as shown by GFAP immunofluorescence. **(F)** CellPhoneDB ligand-receptor interaction map between VEGFA expressed by astrocyte subclusters and VEGFA receptors expressed by all other cell types.

This shared neurodegenerative microglial phenotype across AMD, MS, and AD involved upregulation of multiple genes implicated in studies of neurodegenerative disease risk. These include *APOE*, a key regulator of the transition between homeostatic and neurotoxic states in microglia [*31*] strongly implicated in risk for AD [*32, 33*] and AMD [*34*]; *TYROBP* which encodes the TREM2 ligand DAP12, mutations of which cause Nasu-Hakola disease characterized by a frontal lobe syndrome and AD-like pathology [*35*] and expression of which is upregulated in white matter microglia in MS lesions; *SPP1* (osteopontin), implicated in microglial activation in brains affected by MS [*36*] and AD [*37*]; and *CTSB*, encoding the major protease in lysosomes cathepsin-B, which is upregulated in microglia responding to Aβ plaques in AD [*37*].

Initiation of the pathologic accumulation of extracellular material occurs by different means in these three forms of neurodegeneration. However, the finding that microglial phagocytic, lipid metabolism, and lysosomal activation pathways are upregulated in the early or acute active stage of all three diseases suggests a convergent role for activated human microglia directed towards clearance of the extracellular deposits of debris. The shared signature, however, is not recapitulated in the late or chronic inactive stage clusters across diseases (Supplementary Fig. 7A), implying that the disease-stage specificity is shared across neurodegenerative diseases.

To validate our findings by visualizing microglial signature differences in tissue, we performed simultaneous immunofluorescence for IBA1, a microglia-associated gene, and in situ hybridization for TYROBP and APOE. On sections of human macula, IBA1-positive cells from patients with dry AMD showed enrichment relative to controls for gene transcripts from *TYROBP* and *APOE*, indicating polarization of a subset of microglia towards the neurodegenerative microglial phenotype in early disease (Fig. 3G,H). The average number of puncta identified per IBA1-positive cell for *TYROBP* was 0.28±0.05 in dry AMD vs. 0.02±0.01 for healthy (p < 1e-10; Chi-square test for 0 vs. >0). The average number of puncta identified per IBA1+ microglia for *APOE* was 0.57±0.09 in dry AMD vs. 0.14±0.03 for healthy (p < 1e-08; Chi-square test for 0 vs. >0). These findings validate the expression of a subset of genes associated with the early activation signature in retinas with AMD as compared to controls.

### Astrocyte activation signature identified in dry AMD is shared across the early phase of multiple neurodegenerative diseases

As our initial analysis implicated astrocytes in disease pathogenesis, we performed similar diffusion homology and cross-disease analysis within the astrocyte populations. Using MELD, we identified 4 clusters of astrocytes at a finer granularity of the diffusion condensation hierarchy: cluster 1 was enriched for cells from healthy controls, cluster 2 was enriched for cells from patients with dry AMD, cluster 3 was enriched for cells from patients with neovascular AMD and cluster 4 had equal numbers of cells from all three conditions (Fig. 4A and Supplementary Fig. 4C). Within dry AMD enriched cluster 2, key activation and degeneration associated genes, such as *GFAP*, *VIM* and *B2M* were upregulated when compared with healthy enriched cluster 1 (Fig. 4D), highlighting the need for a cross disease comparison of the astrocytes.

Using MELD, we identified topological features that isolated phase specific populations within MS and AD astrocytes. In both diseases, we identified three clusters: clusters 1 was enriched for cells from healthy brain tissue, clusters 2 was enriched for cells from early pathological AD tissue or acute active MS lesions and clusters 3 was enriched for cells from late pathological AD tissue or chronic inactive MS lesions (Fig. 4B-C). By comparing healthy enriched cluster 1 and early disease enriched cluster 2 within each dataset, we identified a shared gene signature enriched in the early pathology subcluster across all three diseases. The integrated gene signature included markers of activated astrocytes, including *VIM*, *GFAP*, *CRYAB*, and *CD81* [*38, 39*], major histocompatibility complex (MHC) class I (*B2M*) [*40, 41*], iron metabolism (*FTH1* and *FTL*), a water channel component implicated in debris clearance (*AQP4*) [*42*], along with lysosomal activation and lipid and amyloid phagocytosis (*CTSB*, *APOE*). Of interest, many upregulated genes were shared between the microglial and astrocyte early activation signatures, suggesting common glial stress pathways may become activated in response to neurodegeneration. Such states may interact with induction of specific disease related functional polarization of astrocytes by activated microglia [*43*]. Furthermore, this persistent topological feature comparison as performed by diffusion condensation significantly increased our ability to identify differentially expressed genes and develop a common signature (Supplementary Fig. 6B) (see Methods).

Comparison of the overall transcriptomic signature across both healthy-enriched clusters 1 and early disease-enriched clusters 2 revealed that the astrocyte activation signature was distinctly enriched in early pathology (Fig. 4E). Recently, a related disease-associated astrocyte signature has been described in a mouse model of AD [*8*]. We built a composite activation signature and disease-associated astrocyte signature and mapped that onto the clusters, identifying that the early disease enriched clusters displayed higher expression of the disease-associated astrocyte gene signature (Fig. 4F). However, the shared astrocyte signature was not recapitulated in the late or chronic inactive stage clusters across diseases (Supplementary Fig. 7B), implying disease stage specificity that is shared across neurodegenerative diseases.

To validate the signature in tissue, we performed simultaneous GFAP immunofluorescence and RNA in situ hybridization for *B2M*, a component of MHC-I and member of the shared gene signature, on sections of human macula (Fig. 4G). The retinal layers occupied by GFAP-positive astrocytes (inner plexiform layer to inner limiting membrane) contained a higher density of *B2M* transcripts in retina affected by dry AMD relative to control retina (p-value = <1e-03, two-sided Student’s t-test).

### Disease-associated genes drive AMD pathology through glial interactions across disease phases

#### Neurodegeneration-associated genes are expressed in cluster-specific activation signatures and drive signaling interactions

Genome wide association studies (GWAS) have identified many genetic variants associated with neurodegenerative diseases [*44, 45, 46*]. To identify how disease-associated genes may contribute to degeneration, it is critical to identify cell types and, more importantly, cellular activation states that aberrantly express them. We mapped the expression of AMD GWAS-associated genes [*46*] on diffusion condensation identified clusters across multiple levels of granularity, identifying many AMD-associated genes, such as *C3*, *APOE* and *TGFBR1*, to be expressed highly by astrocytes and microglia. We then explored how these specific risk genes map onto the previously identified microglial and astrocytes cellular states (Fig. 5A). Visualizing the expression of the disease-associated risk loci on functional subtypes of glia revealed that many are expressed significantly more by cells in the dry AMD-enriched cluster 2 of both astrocytes and microglia than in other clusters. We performed this analysis for AD and MS disease-specific risk loci across multiple levels of granularity and identified a large set of disease-associated genes expressed highly in early or acute active disease enriched cluster 2 of astrocytes and microglia (Supplementary Fig. 9B).

As many of the risk loci mapped to cluster 2 of astrocytes and microglia play roles in signaling, we used the computational tool CellPhoneDB to identify ligand-receptor interactions between glial disease phase enriched populations [*47*]. Mapping the AMD-associated gene *C3* expressed by astrocyte subpopulations to complement receptors expressed by microglia, the dry AMD-enriched cluster 2 of astrocytes most strongly interacted with the dry AMD-enriched cluster 2 of microglia through complement receptors. Similarly, the AMD-associated gene *APOE*, which is expressed by cluster 2 of dry AMD-enriched microglia most significantly interacted with the APOE-receptors of dry AMD-enriched cluster 2 of astrocytes (Fig. 5B). RNA in situ hybridization of *C3* and *APOE* demonstrated increased expression by astrocytes and microglia respectively in dry AMD retinas, showing that these signaling interactions are upregulated in the early stages of disease (Supplementary Fig. 10). Using DREMI [*48*], we find that the signaling interactions, *C3* on microglia and *APOE* on astrocytes, are strongly associated with our previously identified activation signatures (Fig. 5C), suggesting potential stimulatory receptor-ligand interactions between astrocytes and microglia enriched in dry AMD. Furthermore, these ligand-receptor interaction dynamics were also found in microglia and astrocytes across neurodegenerative conditions, implying that not only are the activation signatures shared but also potentially the mechanisms through which they arise (Supplementary Fig. 9C,D). Given the central role of complement in dry AMD and ongoing complement inhibitor clinical trials [*49*] as well as APOE’s role in plaque accumulation [*50*] and glial cell activation [*51, 52*], these signaling interactions reveal a potential positive feedback loop that may be relevant for therapeutic intervention in all three neurodegenerative conditions. These findings generate testable hypotheses as to the potential molecular signaling interactions between microglia and astrocytes enriched in the early phase of disease that may be relevant for activation of the innate immune response.

#### Astrocyte expression of GWAS-associated gene VEGF drives neovascularization in late AMD

Expression of the AMD GWAS-associated gene *VEGFA* is markedly increased in cluster 3 of astrocytes, a population of cells enriched in late-stage neovascular AMD. When compared to healthy enriched cluster 1, cluster 3 revealed elevation of VEGFA, NR2E1 and HIF1A expression (Fig. 5D), all of which are regulators of cellular responses to low oxygen tension [*53, 54, 55*]. While VEGFA is known to be an important mediator of the neovascularization that characterizes late stage neovascular AMD and the target of therapies for the treatment of disease [*56, 34*], our data provide the first demonstration in humans of a specific subpopulation of retinal astrocytes that are a source of this signal. Simultaneous GFAP immunofluorescence and RNA in situ hybridization for *HIF1A* and *VEGFA* showed increased expression of *HIF1A* throughout the retinal layers and of *VEGFA* in layers with abundant astrocytes in late stage neovascular AMD, with minimal expression in control retinas (Fig. 5E). Finally, using CellPhoneDB interaction analysis [*57*], we identified that VEGFA produced by cluster 3 of astrocytes most strongly interacted with vascular cells, driving pathogenic vessel growth characteristic of late stage, neovascular AMD (Fig. 5F). Thus, while key elements of functional states in specific subsets of glia are shared across the early or active stage of neurodegenerative diseases, diverging functional states specific to disease pathogenicity occur during later stages of disease progression.

## 5 Discussion

Here, we use snRNA-seq to generate the first single-cell transcriptomic atlas of AMD across stages, as well as develop a novel topologically-inspired machine learning approach that allows for meaningful comparison between cell types and states across diseases and phases. Clustering approaches currently applied to single cell data often fail to recognize disease-enriched populations leading to diluted condition-specific signatures. To create rich signatures for cross-disease comparison among rare cellular subpopulations, we created an extended form of diffusion condensation, a novel topologically-inspired approach that learns groups of related cells across granularities. Diffusion condensation combines manifold geometry and topological data analysis frameworks by iteratively applying a low pass filtering process to the intrinsic geometry of single cell data. This approach moves cells progressively towards the center of increasingly global neighborhoods, effectively computing a diffusion homology of the cellular state space. Through persistence analysis, we identify in an unbiased manner, transcriptionally similar cell populations at all levels of granularity. Given healthy and diseased samples of the same tissue, our approach can be applied to the combined data to easily find granularities which maximally isolate disease-enriched cellular populations. This algorithm will be widely useful in more accurately identifying and characterizing cellular populations of interest, both as an unbiased descriptive tool and as an analysis of enrichment in conditions of interest.

Overlaying disease phase enrichment signals onto our multigranular approach, we identified rare subsets of microglia and astrocytes enriched in the early and advanced stages of AMD, AD and MS. Comparing early disease-enriched microglia and astrocyte populations across diseases identified clear activation signatures relating to phagocytic, lipid metabolism, and lysosomal functions. While initial inciting events may differ between degenerative conditions, lipid-rich extracellular plaques play a prominent role in each condition. It is likely that these glial cells are coordinating clearance of extracellular debris and, in turn, becoming activated. While the initial phagocytic clearance can be beneficial, glial activation has been shown to play a role in degeneration in AMD, AD and MS. In later stages of disease, this shared landscape is largely replaced by a disease-specific state, as typified by the proangiogenic astrocyte population that drives pathologic neovascularization in late stage AMD.

In order to link the role of genetic variants identified by population-based GWAS studies to risk of disease, we mapped the expression of disease-associated genes to populations of cells across granularities. This type of multigranular analysis identified not only the cell type but also the activation states through which these genes could influence disease susceptibility. Specifically, we identified key disease-associated genes, *C3* and *APOE* expressed highly by activated populations of astrocytes and microglia respectively, to form a positive feedback signaling loop. This circuit may promote ongoing glial activation in early disease pathogenesis, a phenomenon shared in MS and AD but not present in late stage disease. This analysis revealed that activated glial subsets may be promoting a positive activation loop, which could promote progressive degeneration. In advanced AMD, however, this shared activation signature is replaced with an AMD-specific proangiogenic astrocyte signature that drives pathologic angiogenesis through the expression of the AMD-associated gene *VEGF* and its interaction with vascular cells. These findings indicate that there are commonalities, both in transcriptomic states and cellular circuits, in the early stages of neurodegeneration.

This set of analyses have clear implications on potential therapeutics for AMD and other neurodegenerative diseases. Currently, anti-VEGF therapy is the only intervention approved to treat AMD and is only effective in the most advanced patients. Our unbiased topological analysis not only identified the cell type specificity of *VEGF* expression but also identified pathogenic signaling interactions, which inhibiting provides clinical benefit to patients. Along the same lines, inhibiting interactions between activated glial states could provide therapeutic benefit to AMD patients if applied early in disease pathogenesis. Furthermore, since these mechanisms are shared across MS and AD, it is plausible that these interventions could provide benefit to patients suffering from other neurodegenerative conditions as well. Identifying promising therapeutic candidates to test in neurodegenerative disease clinical trials remains vitally important, and our data suggest that approaches targeting glia may be broadly applicable to the early phases of multiple neurodegenerative diseases.

## 6 Acknowledgements

We would like to thank the retina donors and their families for their contribution to this work. Without their sacrifice our study would not have been possible.

This work was supported by NEI K08-EY026652, the Thome Memorial Foundation (both to B.P.H.), by NIAID training grant 1F30-AI157270 (to M.K.) and by NIAID 5U19-AI089992-08 (to S.K.) and by NIGMS 1RO1-1355929 (to S.K. and G.W.). We thank the Advancing Sight Network and the Lions Gift of Sight Eye Bank for timely retrieval of donor eyes.

## 7 References and Notes

## 8 Supplemental Materials

### 8.1 Materials and Methods

#### 8.1.1 Overview of diffusion Condensation Algorithm

We utilize diffusion condensation as a dynamic process for understanding the diffusion homology of a single cell dataset. Our approach learns data topology by slowly condensing data points towards their neighbors as determined by a diffusion operator. This process is iterative, with each iteration involving the recomputation of a data diffusion operator and further condensation of the data points. A natural byproduct of this process is the emergence of topological features as the condensation proceeds. Normally, mapping of topological features to underlying cellular components is difficult. However, with our agglomerative approach to identifying connected components, diffusion condensation is able to effectively group cells together at all levels of granularity. Persistence analysis of the resultant diffusion homology allows for easy understanding of salient topological features in the underlying cellular manifold. While late-emerging topological features could correspond to large cellular populations such as astrocytes, and bipolar cells, earlier topological features could correspond to smaller populations, such as microglia, vascular cells or subpopulations of larger populations.

In the following sections, we provide a thorough description of each aspect of the diffusion condensation algorithm. This includes a background on topological data analysis, the diffusion condensation process, explanations of algorithm design choices, and information on how comparisons between algorithms were run.

##### Background on topological data analysis

Topological data analysis aims to combine geometric and topological perspectives into a single framework. The underlying idea is that geometry is useful if we are interested in precise measurements and notions of distance between objects, whereas topology is useful if we are interested in describing the relationships between objects. A hybrid perspective can be appealing in situations such as ours, where the individual coordinates of data points are less important than their agglomeration behavior.

Persistent homology refers to a specific topological data analysis framework that is well-equipped to handle geometric data at different granularities. Originally meant to analyze distance functions on sampled manifolds, persistent homology works by approximating the manifold that is underlying a dataset X. This is achieved by constructing a simplicial complex [*58*], i.e., a generalization of a graph, containing higher-dimensional structural elements called simplices, which are subsets of the data points. For a distance threshold *δ*, such a simplicial complex consists of all simplices whose pairwise distances are less than or equal to *δ*, such that

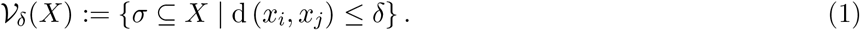

A subset *σ* of cardinality *k* – 1 is referred to as a k-simplex. Thus, the 0-simplices are the vertices of 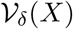, the 1-simplices are the edges, and so on. For *k* ≥ 1, a k-simplex 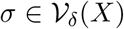is assigned a weight *w_*σ*_* based on the distance of its vertices so that 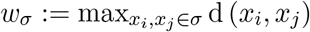. The simplicial complex 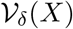 serves as a backbone of the dataset *X*, combining a geometrical perspective (imbued via *δ*) with a topological one: the weighted simplices serve to describe topological features such as connected components (0D), cycles (1D), and voids (2D) in *X*, thus constituting a hybrid between a purely geometrical approach (focusing only on points) and purely topological one (focusing only on connectivity without incorporating distance information). Persistent homology characterizes the evolution of topological features at multiple granularities, as determined by *δ*. For instance, if *δ* is sufficiently large, all points are connected to each other, whereas for *δ* ≈ 0, there will be virtually no connections. Topological features can be efficiently tracked over all potential values of *δ*, and each feature is assigned a *persistence*, a quantity that indicates over which granularities (values of *δ*) a feature is present. For example, if *X* is a densely-sampled square, it will intrinsically have one connected component, which will be assigned a high persistence. Information about the persistence of features is collected in topological descriptors, such as persistence diagrams [*59*] or persistence barcodes [*22*]. In a persistence barcode, each topological feature is represented as a horizontal bar whose length indicates its persistence. Topological features that persist longer (i.e., have longer bars) are considered to be more unique, while features that persist for fewer iterations are considered ubiquitous or even noisy. The barcode metaphor yields a powerful visualization, which serves as an informative summary of a dataset’s underlying features. Until now, topological frameworks and analysis of persistent homology has not been applied to single cell analysis.

##### Prior work in multigranular data abstraction

Previous attempts at providing multigranular data abstraction of single cell data rely on hierarchical clustering, a family of methods that attempts to derive a tree of clusters based on either recursive recursive splitting or agglomeration of data points. Splitting based approaches, including recursive bisection [*60*] and divisive analysis clustering [*61*], work in an iterative, top-down fashion, each time optimizing a partition of the data into clusters. Agglomerative methods meanwhile, including the popular linkage clustering, or community detection methods such as Louvain [*13*], work in a bottom-up fashion recursively merging points into clusters. While intuitively more related to the merges in diffusion condensation, there is a fundamental difference between the coarse graining operation applied here and the greedy agglomeration approaches: hierarchical methods recursively force merges or splits in data. Although a hierarchy of cells is created through this approach, concepts of data-driven topology such as persistence calculations cannot readily be applied. Since the merging of points is arbitrary and not scaled by differences between underlying cells, the persistence of a population largely does not have meaning, creating an incomplete understanding of the topology of the data.

Inspired by concepts of persistence homology and dearth of current computational techniques amenable to such analysis, we decided to create a topologically-inspired approach to understand the multigranular structure of single cells based on their inherent manifold geometry.

##### Background in manifold learning and diffusion filters

High dimensional data can often be modeled as originating from a sampling 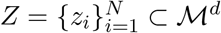 of a d-dimensional manifold 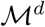 that is mapped to observations of dimension *n* ≫ *d*, collected in *X* = {*x*_1_,…, *x_N_*} ⊂ ℝ^*n*^ via a nonlinear function *x_i_* = *f(z_i_*). Intuitively, the reason for this phenomenon is that data collection measurements (modeled here via *f*) typically result in high dimensional observations, even when the intrinsic dimensionality, or degrees of freedom, in the data is relatively low. This manifold assumption is at the core of the vast field of manifold learning (e.g., [*62, 63, 64, 65*], and references therein), which leverages the intrinsic data geometry, modeled as a manifold, for exploring and understanding patterns, trends, and structure in data.

In [*63*], diffusion maps were proposed as a robust way to capture intrinsic manifold geometry in data using random walks that aggregate local affinity to reveal nonlinear relations in data and allow their embedding in low dimensional coordinates. These local affinities are commonly constructed using a Gaussian kernel

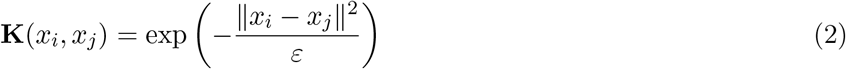

for *i, j* ∈ {1,…, *N*}, where **K** is an *N* × *N* Gram matrix whose (*i, j*) entry is denoted by **K**(*x_i_*, *x_j_*) to emphasize the dependency on the data *X*. The bandwidth parameter *ε* controls neighborhood sizes. A diffusion operator is defined as the row-stochastic matrix **P** = **D**^−1^**K** where **D** is a diagonal matrix with **D**(*x_i_*, *x_i_*)= ∑_*j*_ **K**(*x_i_*, *x_j_*), which is referred to as the degree of *x_i_*. The matrix **P** defines single-step transition probabilities for a time-homogeneous diffusion process (which is a Markovian random walk) over the data, and is thus referred to as the diffusion operator. Furthermore, as shown in [*63*], powers of this matrix **P**^*t*^, for *t* > 0, can be used for multiscale organization of *X*, which can be interpreted geometrically when the manifold assumption is satisfied.

While originally conceived for dimensionality reduction via the eigendecomposition of the matrix **P**, recent works [*66, 67, 68, 69*] have extended the diffusion framework of [*63*] to allow processing of data features by direct application of the operator **P**. These approaches include data denoising and imputation [*66*], data generation [*67*], and graph embedding with geometric scattering [*68, 69*]. In these cases, **P** serves as a smoothing operator, and may be regarded as a generalization of a low-pass filter for either unstructured or graph-structured data. Indeed, consider a vector **v** ∈ ℝ^*N*^ that we think of as a signal **v**(*x_i_*) over X. Then **Pv**(*x_i_*) replaces the value **v**(*x_i_*) with a weighted average of the values **v**(*x_j_*) for those points *x_j_* such that 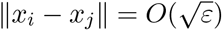.

Each of these applications, however, uses a time homogeneous matrix **P** the defines the transition probabilities of a random walk over the data set *X*. Computing powers of **P** runs the walk forward, so that **P**^*t*^ gives the transition probabilities of the t-step random walk. Since the same transition probabilities are used for every step of the walk, the resulting diffusion process is time homogeneous.

By contrast, a time inhomogeneous diffusion process arises from an inhomogeneous random walk in which the transition probabilities change with every step. Its t-step transition probabilities are given by

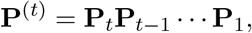

where **P**^(*t*)^ is a time inhomogeneous diffusion operator is composed of many **P**_*k*_ is the Markov matrices that encodes the transition probabilities at step k and **P**^*t*^ describes transition probabilities for the time homogeneous case (where the resulting matrix is a power of the diffusion operator **P**).

Previous works have applied the time inhomogeneous diffusion operator to data with an explicit time variable [*70*], attempting to approximates heat diffusion over the time varying manifold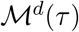. The application of time inhomogeneous or homogeneous diffusion operators to learn data topology however, has not been previously explored.

##### Overview of diffusion condensation

Diffusion condensation is a dynamic process the builds upon previously established concepts in diffusion filters, diffusion geometry and topological data analysis. The algorithm slowly and iteratively moves points together at a rate determined by the diffusion probabilities between them [*10*]. This iterative process reveals the topology of the underlying geometry and identifies key persistent features that can be directly mapped back onto the underlying dataset. The diffusion condensation approach involves two steps that are iteratively repeated until all points converge:

1. Compute a time inhomogeneous Markov diffusion operator from the data
2. Apply this operator to the data as a low-pass diffusion filter, moving points towards local centers of gravity

As established in prior work, the application of the operator **P** to a vector **v** averages the values of **v** over small neighborhoods in the data [*66*]. In the case of data *X* = {*x*_1_,…, *x_N_*}⊂ ℝ^*n*^ measured from an underlying manifold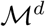 with the model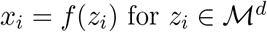, this averaging operator can be directly applied to the coordinate functions *f* = (*f*_1_,…, *f_n_*). Let **f**_*k*_ ∈ ℝ^*N*^ be the vector corresponding to the coordinate function *f_k_* evaluated on the data samples, i.e., **f**_*k*_(*z_i_*)= *f_k_*(*z_i_*). The resulting description of the data is given by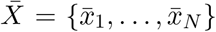 where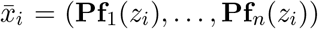. Applications of the operator **P** to *X* dampens high frequency variations in the coordinate function and creates smoothed output 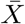.

Here we consider not only the task of eliminating variability that originates from noise, but also coarse graining the data coordinates to learn the topology of the data. Therefore, we aim to gradually eliminate local variability in the data using a time inhomogeneous diffusion process that refines the constructed geometry to coarser resolutions as time progresses. This condensation process proceeds as follows. Let *X*(0) = *X* be the original data set with Markov matrix **P**_0_ = **P** and 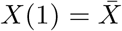 the coordinate-smoothed data described in the previous paragraph. We can iterate this process to further reduce the variability in the data by computing the Markov matrix **P**_1_ using the coordinate representation *X*(1). A new coordinate representation *X*(2) is obtained by applying **P**_1_ to the coordinate functions of *X*(1). In general, one can apply the process for an arbitrary number of steps, which results in the condensation process. Let *X*(*t*) be the coordinate representation of the data after *t* ≥ 0 steps so that *X*(*t*)= {*x*_1_(*t*),…, *x_N_* (*t*)} with 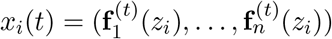, where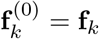. We obtain *X*(t + 1) by applying **P**_*t*_, the Markov matrix computed from *X*(*t*), to the coordinate vectors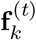. For *t* ≥ 0, this process results in

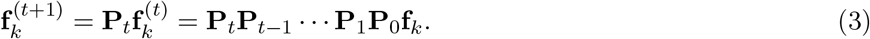

From (3), we see that the coordinate functions of the condensation process at time *t* +1 are derived from the imposed time inhomogeneous diffusion process **P**^(*t*)^ = **P**_*t*_… **P**_0_. The low-pass operator **P**_*t*_ applies a localized smoothing operation to the coordinate functions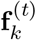. Over the entire condensation time, however, the original coordinate functions **f**_*k*_ are smoothed by the cascade of diffusion operators **P**_*t*_…**P**_0_. This process adaptively removes the high frequency variations in the original coordinate functions. The effect on the data points *X* is to draw them towards local barycenters, which are defined by the inhomogeneous diffusion process. Once two or more points collapse into the same barycenter, they are identified as being members of the same topological feature.

##### Adaptions to diffusion condensation for application to single-cell RNA-sequencing data

In its original form, the diffusion condensation process does not easily apply to scRNAseq data as it does not condense points on the biological manifold and does not scale to hundreds of thousands of cells. To address these concerns, we have made the following three significant adaptions for application to single cell data:

1. transforming cells to a novel manifold coordinate system,
2. merging points together that fall below a preset distance threshold *ζ* to increase computational efficiency,
3. identifying persistent topological features by analyzing diffusion homology

Each of these steps is explained below.

##### Novel manifold coordinate system

The original diffusion condensation algorithm [*10*] implements the coarse grainings of the data in the ambient measurement space. However, condensing in this space can lead to “averaged” points that deviate from the intrinsic data manifold, especially in cases where the intrinsic manifold is very curved. As cellular state spaces can be heavily non-linear [*18, 48, 21*], we require an alternative method of diffusion condensation that ensures that the condensed points remain on the manifold. Rather than using the original features, we use the potential representation of data points used in PHATE [*18*] to learn the geometry of the manifold and allow data points to condense on the manifold and not in non-linear measurement space.

##### Data merging

When two or more points collapse into the same barycenter, as defined by a preset distance threshold *ζ*, we merge them into a single topological feature since they would then have approximately the same manifold coordinates. This has the effect of density subsampling the data iteratively, and allowing for subsequent iterations to proceed faster as the matrix has reduced dimensionality. After this merging operation, we treat the topological feature as a single point and in our diffusion homology analysis, trace its persistence across iterations. As we iterate this process, the condensation slowly coarse grains the data to reveal structure at all levels of granularity while avoiding the typical tendency of traditional hierarchical clustering approaches to force cluster merges at every scale. This adaption greatly increases the speed of condensation when applied to large biological datasets and prevents smaller cellular populations from being absorbed into more abundant populations.

We can also describe this data merging through the lens of *diffusion homology*, a novel technique inspired by persistent homology, which describes the evolution of 0-dimensional topological features—connected components— during the condensation process. Specifically, we start with every point *x_i_* ∈ *X* in its own connected component. As the diffusion condensation iteration *t* increases, points are moving closer towards local barycenters. If the distance of points *x_i_* and *x_j_* falls below a preset distance threshold *ζ* we merge them together. In topological terms, this has the effect of merging two connected components, the one created by *x_i_* and the one created by *x_j_*. If such a merge happens in step *t* of the diffusion condensation process and the two points are not already part of the same connected component, we associate a persistence of *t* with the connected component that gets merged. Furthermore, we visualize all such merges as a barcode diagram to summarize the diffusion condensation process (Fig. 1D). Following the discussion of topological data analysis above, such merges can be considered akin to a persistent homology calculation. The crucial difference is that we are not analyzing topological features for varying values of *δ*, the distance threshold for the simplicial complex approximation. Instead, we are tracking changes as the underlying dataset *X* is being updated in consecutive iterations of the diffusion condensation process. We therefore refer to the process of tracking topological features along this process as *diffusion homology*.

##### Persistence-inspired analysis of diffusion homology

Diffusion homology can be calculated efficiently by using a disjoint set (or union–find) data structure [*71*]. This data structure keeps track of the data points associated with a specific connected component, offering the operations FIND to query the component a point belongs to (initially, `FIND`(*x_i_*)= *x_i_*, as described above), and `UNION`(*x_i_*, *x_j_*) to merge the two connected components *x_i_* and *x_j_* belong to, respectively, into one. Specifically, for all pairs *x_i_*, *x_j_* with d(*x_i_*, *x_j_*) ≤ *ζ* in iteration *t*, we perform `UNION`(*x_i_*, *x_j_*) if and only if `FIND`(*x_i_*) ≠ `FIND`(*x_j_*), thus ensuring that only merges between points belonging to *different* connected components are performed in each step. Diffusion homology is easy to calculate, as the disjoint set data structure for *n* data points can be implemented to have a worst-case complexity of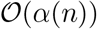, where *α*(n) is the inverse Ackermann function, which is extremely slow growing. For all intents and purposes, the operation therefore have a maximum complexity that is linear in the number of points. Since finding candidate points to merge requires *at least* one pass through the whole dataset in the worst case, diffusion homology calculations do not impair computational performance of the diffusion condensation process.

The resulting diffusion homology barcode is a useful summary of the diffusion condensation process. Longer bars correspond to structures that are more prominent because they occur over different granularities of the diffusion condensation. The occurrence of such prominent barcodes indicates that the corresponding components in the data are all strongly connected to one another and not part of less prominent connected components, represented by shorter bars. In our single cell analysis, these persistent topological features indicate that the underlying cells which create this feature are related closely to one another at coarse granularity. These cells likely can be accurately be called with a cell type label such as astrocytes or bipolar cells. Short bars, by contrast, represent small subsets of points that get merged early on and quickly are subsumed by more persistent connected components. The barcode can therefore not only be used to assess potential granularities at which to view a dataset but also identify components that are heavily interconnected with one another at a particular granularity.

##### Combining diffusion condensation with MELD to identify populations enriched in disease phases

While diffusion condensation and diffusion homology analysis may be adept at identifying topological features, identifying granularities that isolate disease enriched populations remains a problem. In order to identify groups of cells enriched in specific disease states, we combined our topological approach with MELD [*23*]. MELD creates a joint graph of the samples being compared, and returns a relative likelihood that quantifies the probability that each cell state in the graph is more likely in a particular disease condition. This likelihood score is found by first computing a cell-cell graph before creating an indicator signal for each of the two conditions. In our work, we use the disease phase of origin as the indicator signal, labeling the cells not from that phase as a control. Then MELD smooths, or low-pass filters these signals over the cell-cell graph using the heat kernel to calculate the relative likelihood of each condition over the cellular manifold. The density estimates of both conditions are then inverted via Bayes formula to create a relative likelihood of the condition given the cell. This likelihood score highlights regions of the manifold enriched in different conditions. Using the MELD likelihood scores, we can identify cellular subsets enriched in particular disease phases and furthermore, identify topological features that maximally separate disease-phase signals.

##### Comparison to clustering approaches applied to single cell analysis

In order to quantify diffusion condensation’s ability to identify cell types and cell states, we created a single cell dataset in which cell types were known at multiple levels of the cellular hierarchy. We downloaded 27,181 cells measured with single-cell RNA-seq (scRNAseq) and CITE-seq from human peripheral blood mononuclear cells (PBMCs) [*12*] and identified cell type labels by gating on known CITE-seq features as is standard in flow cytometry [*72*]. Cell type labels were identified in a hierarchical manner. At the most coarse granularity, lineages were identified: myeloid, dendritic and lymphocytic. Each of these were then broken down into cell types, i.e. lymphocyte lineage was broken into B, T and NK cells. Finally, each of these cell types were broken down into subtype, i.e. T cells were broken into CD4^+^ and CD8^+^ T cells. With these labels serving as ground truth, clusters were identified on the scRNAseq data using Louvain, DBscan, agglomerative clustering and diffusion condensation. As each of these algorithms create clusters across granularities, we compared all clustering granularities produced from each algorithm with each of our ground truth hierarchical cell type labels using Adjusted Rand Index (ARI). Across all comparisons, diffusion condensation more optimally identified ground truth cluster labels when compared to Louvain, DBscan and agglomerative clustering. This finding indicates that diffusion condensations topological approach optimally identifies cellular populations and subpopulations all throughout its diffusion homology.

##### Topological analysis improves ability to find shared gene signatures

Typically, disease associated signatures are uncovered by performing differential expression analysis between cells from the disease condition and cells from the healthy condition at the resolution of cell type. For instance, microglia would be separated into two groups based on condition of origin, either disease or healthy, which would then be compared. We believe that comparing cellular states identified with topological data analysis enriched for different conditions of origin will identify transcriptomic differences and similarities more rigorously.

To illustrate this point, we performed differential expression analysis between microglia based on their condition of origin across all three diseases. After setting significance a cutoff established in our topological analysis (EMD cutoff of 0.044, 0.130 and 0.164 for AMD, MS and AD respectively), we identified significantly enriched genes in the early or acute active phase of each disease (Supplementary Fig. 6A). However, across all comparisons, we identified significantly fewer differentially expressed genes in this cell type analysis (135, 68 and 416) than in the topological analysis (618, 795 and 1551 for AMD, AD and MS respectively), indicating that cellular subtyping with diffusion condensation before comparison increases our ability to detect differentially expressed genes. In cross-disease comparisons among early stage neurodegenerative microglia, only 17 common genes were found, significantly less than the 168 common genes found with topological analysis. Of the common genes, only half of the activation signature was found (*APOE*, *B2M*, *FTH1*, *FTL*, *SPP1*).

Similar to our coarse grained microglial comparison, we compared the strength of our topological approach in astrocytes. Comparing astrocyte states based on condition of origin with previously set significance cut offs (0.034, 0.061 and 0.071 for AMD, AD and MS respectively), we identified significantly fewer enriched genes (221, 271 and 886) than we found with our topological analysis (1444, 680 and 2278 genes for AMD, AD and MS respectively) (Supplementary Fig. 6B). In our cell type level analysis, only 28 common genes were found, significantly less than the 630 common genes found with topological approach. Of the common genes, only half of the activation signature was found (*AQP4*, *CD81*, *CRYAB*, *GFAP*).

Collectively, these comparisons reveal the strength of finer grained analysis with diffusion condensation.

##### Software Availability

The Diffusion Condensation package, as implemented in python, is available for download with a guided tutorial on the Krishnaswamy Lab Github page: https://github.com/KrishnaswamyLab/Diffusion_Condensation.

#### 8.1.2 Other of Computational Methods

##### Single-nucleus AMD RNA sequencing pre-processing

snRNA-seq data from macular samples, were processed according to the following steps. Sample demultiplexing and read alignment to the NCBI reference pre-mRNA GRCh38 was completed to map reads to both unspliced pre-mRNA and mature mRNA transcripts using CellRanger version 3.1.0. Gene and cell matrices from the remaining retinas with dry AMD (n=3), neovascular AMD (n=4), and healthy controls (n=6) were combined into a single file. We prefiltered using parameters in scprep (v1.0.3, https://github.com/KrishnaswamyLab/scprep). Cells that contained at least 1400 unique transcripts were kept for further analysis to generate a cell by gene matrix containing 50,498 cells. Normalization was performed using default parameters with L1 normalization, adjusting total library side of each cell to 1000. Any cell with greater than 200 normalized counts of mitochondrial mRNA was removed. Batch correction was performed using Harmony (https://github.com/immunogenomics/harmony) to align batch effects introduced by sequencing batch, postmortem interval, sample acquisition location and 10X sequencing chemistry [*73*]. Raw and processed data files for human snRNA-seq data will be available for download through GEO under an accession number to be assigned with no restrictions on data availability.

##### Single-nucleus AD and MS RNA sequencing pre-processing

snRNA-seq data for AD and MS was acquired from published sources [*6, 30*]. Cells that contained at least 1000 unique transcripts were kept for further analysis to generate a cell by gene matrix for each disease. Normalization was performed using scprep default parameters with L1 normalization, adjusting total library side of each cell to 1000. Any cell with greater than 200 normalized counts of mitochondrial mRNA was removed. Batch correction was performed on MS data using Harmony (https://github.com/immunogenomics/harmony) to align batch effects introduced by sequencing batch, capture batch and sex.

##### Cell type identification with diffusion condensation

All cell types were identified by performing persistence analysis on the diffusion condensation calculated diffusion homology. In order to identify cell types, we calculated the most persistent topological features in each dataset and assigned them to a cell type based on cell type specific marker genes. Often, a topological feature expressed marker genes from more than one cell type, in which case we asked for less persistent topological features within that feature. This process continued until all cell types were identified. Distinguishing microglia from macrophages in the CNS during neuroinflammation can be challenging as hematopoietic stem cell-derived macrophages can enter the CNS and dynamically change their transcriptional profile to adopt a microglial genetic signature. Thus we cannot exclude hematopoietic stem cell-derived macrophages from the analyses [*74*].

##### Cell type specific abundance analysis

In order to determine the cell types most enriched in disease, we performed cell type specific abundance analysis. After normalizing for overall condition specific abundance differences, we would expect half of the cells of any cell type to arise from the disease condition and half to arise from the healthy condition. Cell types which deviated significantly from this expectation indicate an enrichment in disease as computed by a single sided binomial test.

##### Differential Expression Analysis

In order to perform differential expression analysis, first each dataset was de-noised with MAGIC [*21*]. When comparing specific subpopulations, differential expression analysis was performed by first computing Earth Mover’s Distance (EMD) between every de-noised gene in each population as previously described [*21*], effectively ranking the degree of differential expression for every gene. For each comparison, the top 10% of EMD values were set as significant.

##### Interaction Analysis

Cell–cell ligand-receptor analysis was conducted on pre-processed snRNA expression data using the CellPhoneDB python package (https://github.com/Teichlab/cellphonedb, v2.1.4) [*57*]. Before conducting analysis, the package database of 834 curated ligand-receptor combinations and multi-unit protein complexes was supplemented with 2557 ligand-receptor interactions found in the celltalker database (https://github.com/arc85/celltalker) [*75*]. The in-built database-generate function was utilized to update the existing database. Our comprehensive user-generated database was invoked in each run of the CellPhoneDB statistical-analysis command function.

Two separate CellPhoneDB interaction maps were computed on differing inputs. First, disease phase enriched microglia and astrocytes with subcluster identity were run to identify signaling interactions between astrocyte and microglial activation states (Fig. 5B). The number of permutations was set to 2,000 and p-value threshold was set to 0.01. The second interaction map was computed between subclusters of astrocytes and all non-astrocyte cell types to identify cell types that interact with astrocyte expressed VEGFA (Fig. 5F). Since all 50,498 cells from the AMD data set were used in this run, we set the number of permutations to 10,000 and p-value threshold to 0.01.

#### 8.1.3 Overview of Biological Methods

##### Human tissues

Postmortem eyes for the Chromium Single Cell 3’ assay (n = 13) and medical records containing AMD disease stage were obtained from Advancing Sight Network (Alabama), Lions Gift of Sight Eye Bank (Minnesota) or the Yale Department of Pathology with a maximum post-mortem interval of 10 hours. Globes were examined for retinal disease by an ophthalmologist (B.P.H.) prior to dissection and dissociation of the samples. This study was approved by the Yale Institutional Review Board. We complied with all relevant ethical regulations for work with human participants, and all human tissue samples were obtained with proper consent prior to tissue collection. Retina for snRNA-seq was obtained from 13 unrelated human post-mortem donors that included normal, dry, and neovascular AMD stages (Supplemental Table 1). For each sample we profiled the macula, which is the region of the retina responsible for central vision and affected by AMD pathology. We selected three samples with drusen, a pathologic sign associated with the dry stage of the disease, and four samples with neovascularization in the advanced stage of the disease. Normal donors had no history of retinal disease. Retinal dissection: Globes were placed in RNAlater (ThermoFisher) and transported on ice. Trephine punches (6 mm diameter) were used to isolate samples from the macula in the central retina, located away from the optic disc and major arterioles. For each punch of tissue, the retina was mechanically separated from the underlying retinal pigment epithelium-choroid, snap-frozen on dry ice and stored at −80°C.

##### Isolation of nuclei from frozen retinal tissue

Nuclei were isolated and purified using the Nuclei EZ Prep Nuclei Isolation Kit (Sigma), following the manufacturer’s protocol, with some modification. All procedures were carried out on ice or at 4°C. Briefly, frozen retinal tissue was subjected to dounce homogenization (25 times with pestle A followed by 25 times with tight pestle B) using the KIMBLE Dounce Tissue Grinder Set (Sigma) in 2ml EZ Lysis buffer. The sample was transferred to a 15ml tube with an additional 2ml EZ lysis buffer and incubated on ice for 5min. Following incubation, the sample was centrifuged at 500xg, 5min at 4°C. Supernatants were discarded, and the isolated nuclei were resuspended in 4ml EZ lysis buffer, incubated for 5min on ice and centrifuged at 500xg for 5min at 4°C. Next, the nuclei were washed with 4ml ice-cold Nuclei Suspension Buffer (1X PBS containing 0.01% BSA and 0.1% RNAse inhibitor), resuspended in 1ml Nuclei EZ Storage buffer and passed through a 40μM nylon cell strainer. The nuclei suspensions were counted with trypan blue prior to loading on the microfluidics platform.

##### Droplet-based microfluids snRNA-seq

Isolated nuclei from each macular sample was processed through microfluidics-based single nuclear RNA-seq. Single-cell libraries were prepared using the Chromium 3’ v2 and v3 platforms (10X Genomics) following the manufacturer’s protocol. Briefly, single nuclei were partitioned into Gel beads in Emulsion in the 10X Chromium Controller instrument followed by lysis and barcoded reverse transcription of RNA, amplification, shearing and 5’ adapter and sample index attachment. On average, 7000 nuclei were loaded on each channel that resulted in the recovery of 4000 nuclei. Libraries were sequenced on the Illumina NextSeq 500 platform. After quality control preprocessing, a total of 50,498 snRNA-seq profiles were used in subsequent analyses. This dataset was corrected for batch effects across samples using the Harmony algorithm [*76*].

##### In situ RNA hybridization and immunofluorescence

To validate the gene expression differences, in situ hybridization was performed using RNAscope Multiplex Fluorescent V2 Assay (Advanced Cell Diagnostics, Hayward, CA, USA). Macula dissected from whole human globes were fixed in 4% paraformaldehyde (PFA) at 4°C overnight. Tissues were sequentially dehydrated with 15% sucrose, then 30% sucrose before embedding in OCT, and frozen on dry ice. OCT molds were sectioned at 10 μmm thickness. RNA in situ hybridization was performed according to the manufacturer’s protocol. Briefly, fixed frozen sections were baked at 60°C for 1 hr prior to incubation in 4% PFA for 10 mins and protease digestion pretreatment. Target probes were hybridized to an HRP-based temperature sensitive signal amplification system, followed by color development. Housekeeping genes POLR2A, PPIB, and UBC were used as internal-control mRNA (Supplementary Fig. 8); if probes for these mRNAs were not visualized, the sample was regarded as not available for gene expression study. The slides were counterstained with DAPI during immunofluorescence protocol (see below). Positive staining was determined by fluorescent punctate dots in the appropriate channels in the nucleus and/or cytoplasm. Following RNA in situ hybridization protocol, fixed frozen sections were blocked with animal serum and incubated overnight at 4°C with primary antibodies (see antibody segment below). Secondary antibody incubation was for 1 hr at room temperature and cell nuclei were counterstained with DAPI. Images were captured immediately using a confocal microscope (Zeiss LSM800, Jena, Germany). The following antibodies against human antigens were used: GFAP (1:500, MA5-12023, Invitrogen) and Iba1 (1:500, 019-19741, Fujifilm). Antibodies were visualized with Alexa Fluor 488 (1:200, A-11001 / A-21208, Invitrogen).

## 8.2 Supplementary Figures

**Supplementary Figure 1:**
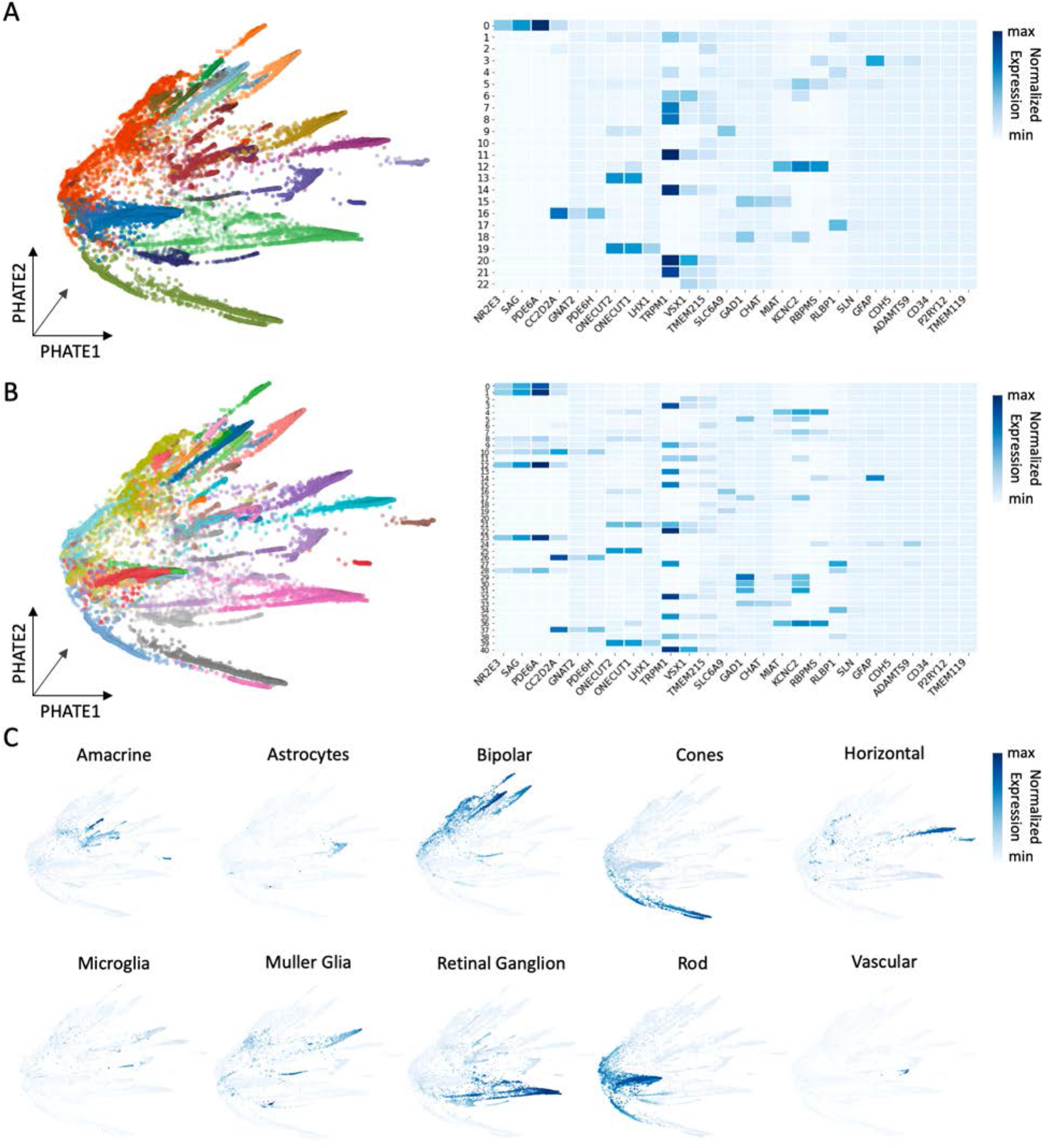
Louvain clustering of AMD single cell data does not identify rare retinal populations across granularities. **(A)** Visualization of 22 coarse grain clusters identified by Louvain. Louvain identified populations are not able to reliably identify cell types, as shown by the average normalized expression of known cell type specific marker genes. **(B)** Visualization of 40 fine grain clusters identified by Louvain. Louvain identified populations are not able to reliably identify cell types, as shown by the average normalized expression of known cell type specific marker genes. **(C)** Cell type specific signatures based on composite normalized expression of cell type specific marker genes visualized on PHATE.

**Supplementary Figure 2:**
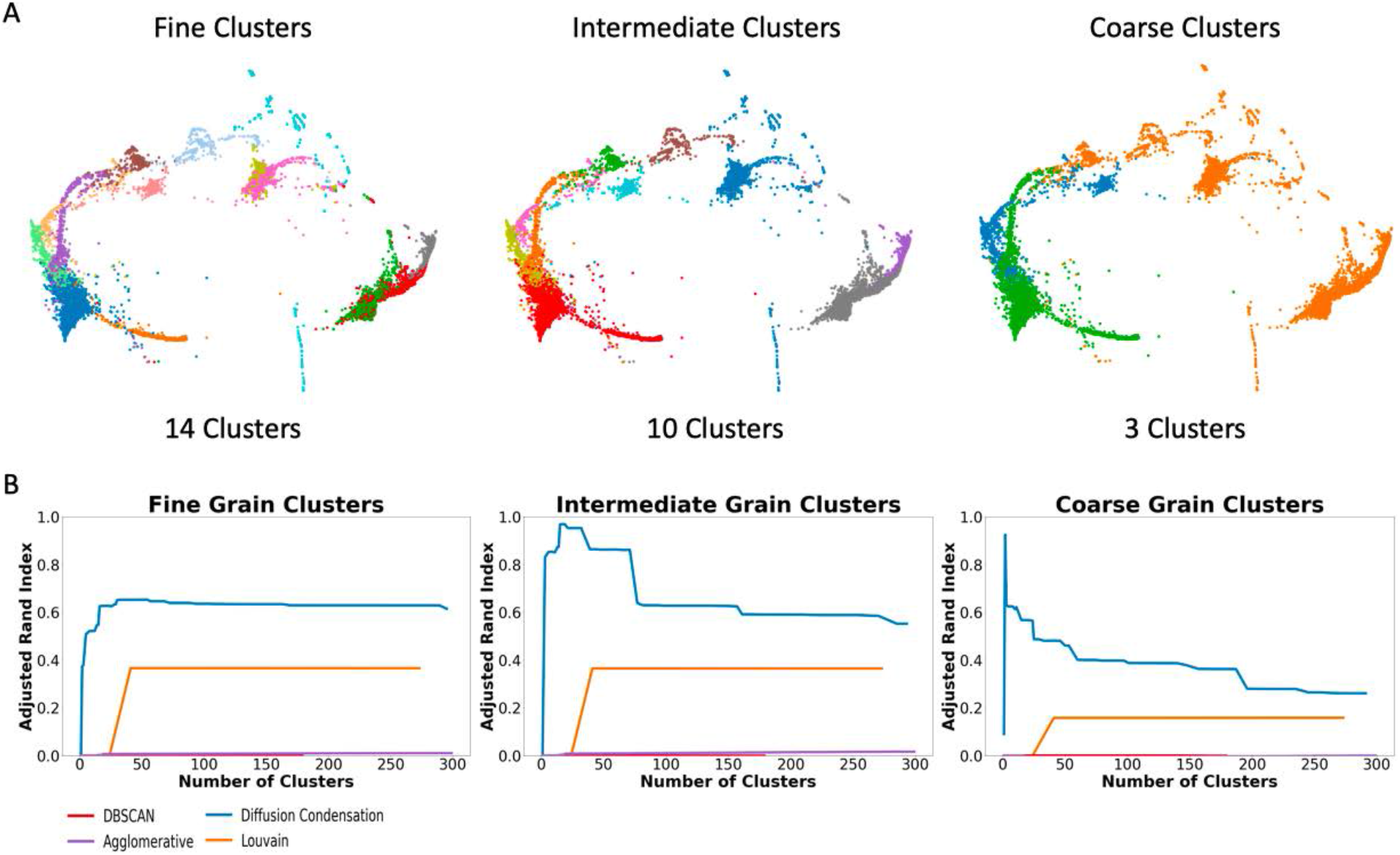
Comparison across granularities between different clustering tools applied to single cell analysis. **(A)**27,181 cells measured with single-cell RNA-seq and CITE-seq from human peripheral blood mononuclear cells (PBMCs) [*12*]. Cell type labels were identified by gating on known CITE-seq features. PBMCs a known hierarchical organization, where identified cell types can be organized at multiple levels of granularity: cellular subset (14 cluster), cell type (10 clusters) and lineages (3 clusters). These identified clusters formed ground truth labels for comparison between clustering approaches. **(B)** Comparison between different clustering approaches that produce multigranular groupings of single cell data: Louvain, agglomerative, DBscan and Diffusion Condensation. Clusters produced from each approach were compared to ground truth cluster labels at each granularity using Adjusted Rand Index (ARI).

**Supplementary Figure 3:**
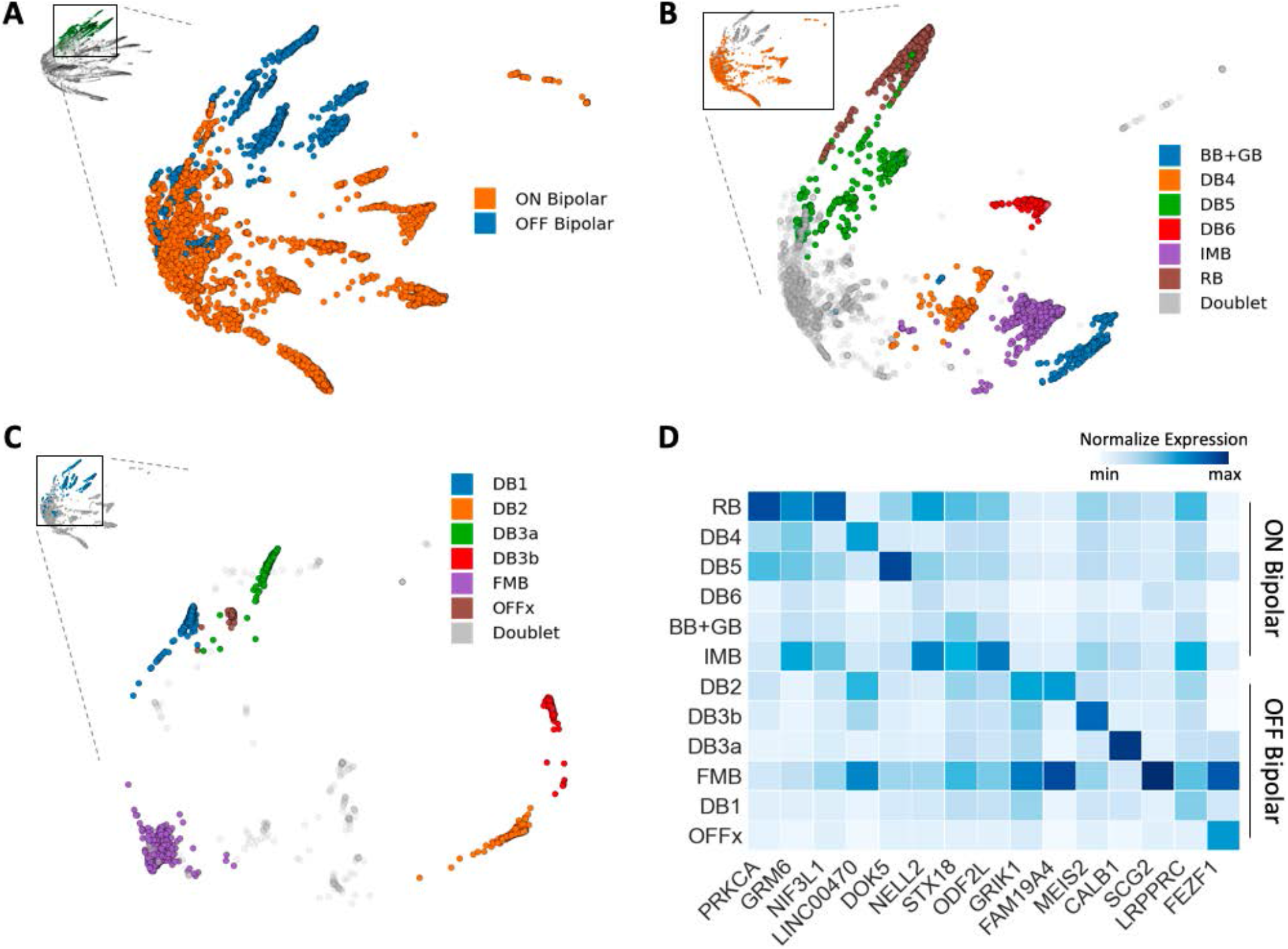
Diffusion condensation identifies known subtypes of bipolar cells across multiple levels of granularity. **(A)** Diffusion condensation identifies ON and OFF bipolar cell subsets with persistence analysis of the diffusion homology. **(B)** Further persistence based analysis of ON bipolar cells reveals known subsets. **(C)** Further persistence based analysis of OFF bipolar cells reveals known subsets. **(D)** Diffusion condensation reliably identifies established cell types, as shown by average normalized expression of known bipolar subset-specific marker genes.

**Supplementary Figure 4:**
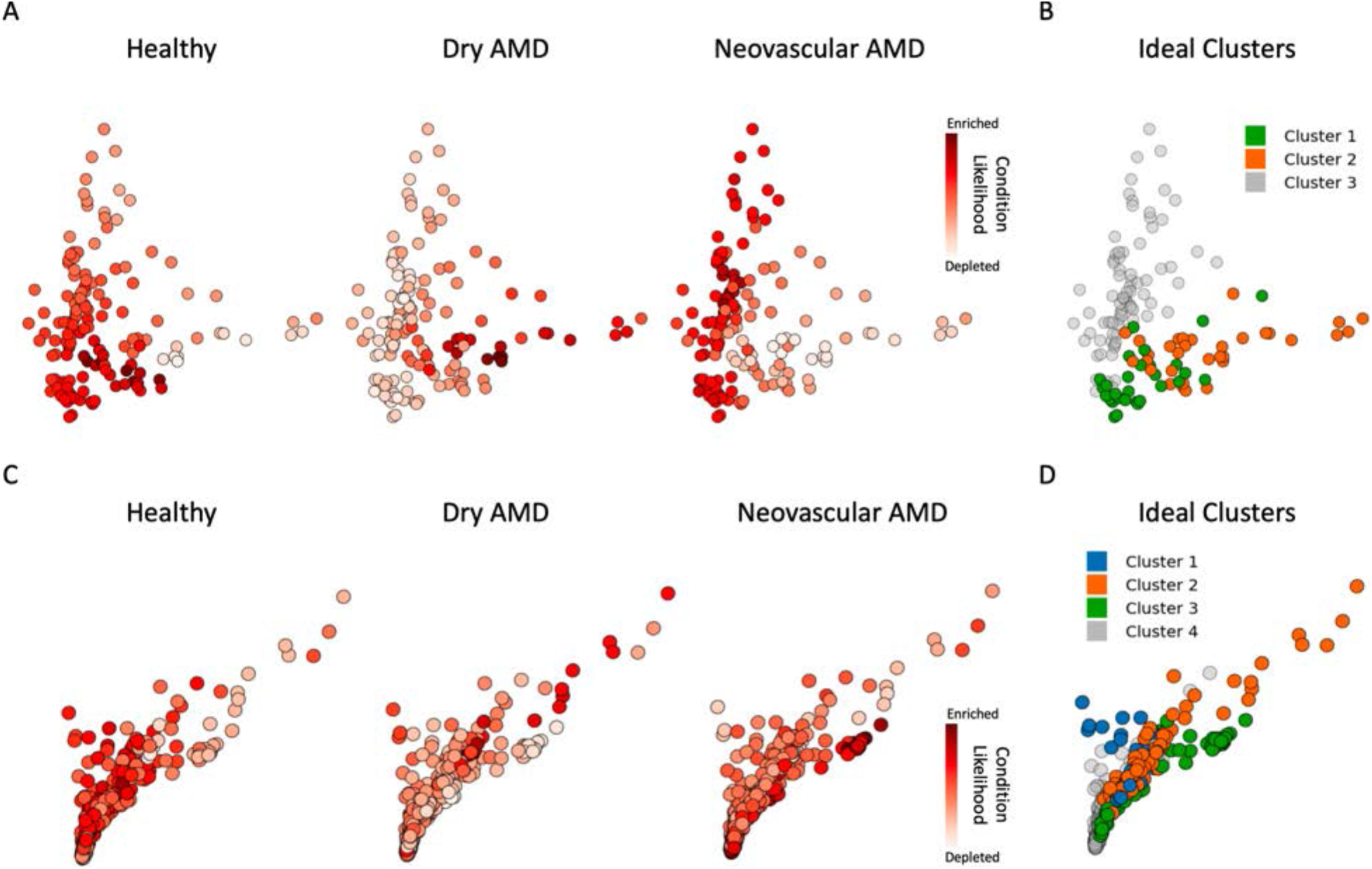
MELD and diffusion condensation analysis of microglia and astrocytes identifies cellular states enriched in healthy, dry AMD, and neovascular AMD. **(A)** MELD likelihood scores identify regions of microglial sub-manifold enriched for cells from healthy retinas, dry AMD retinas and neovascular AMD retinas. **(B)** Diffusion condensation identifies three topological features which isolated each of the microglia MELD condition likelihood scores. **(C)** MELD likelihood scores identify regions of astrocyte sub-manifold enriched for cells from healthy retinas, dry AMD retinas and neovascular AMD retinas. **(D)** Diffusion condensation identifies four topological features which isolated each of the astrocyte MELD condition likelihood scores.

**Supplementary Figure 5:**
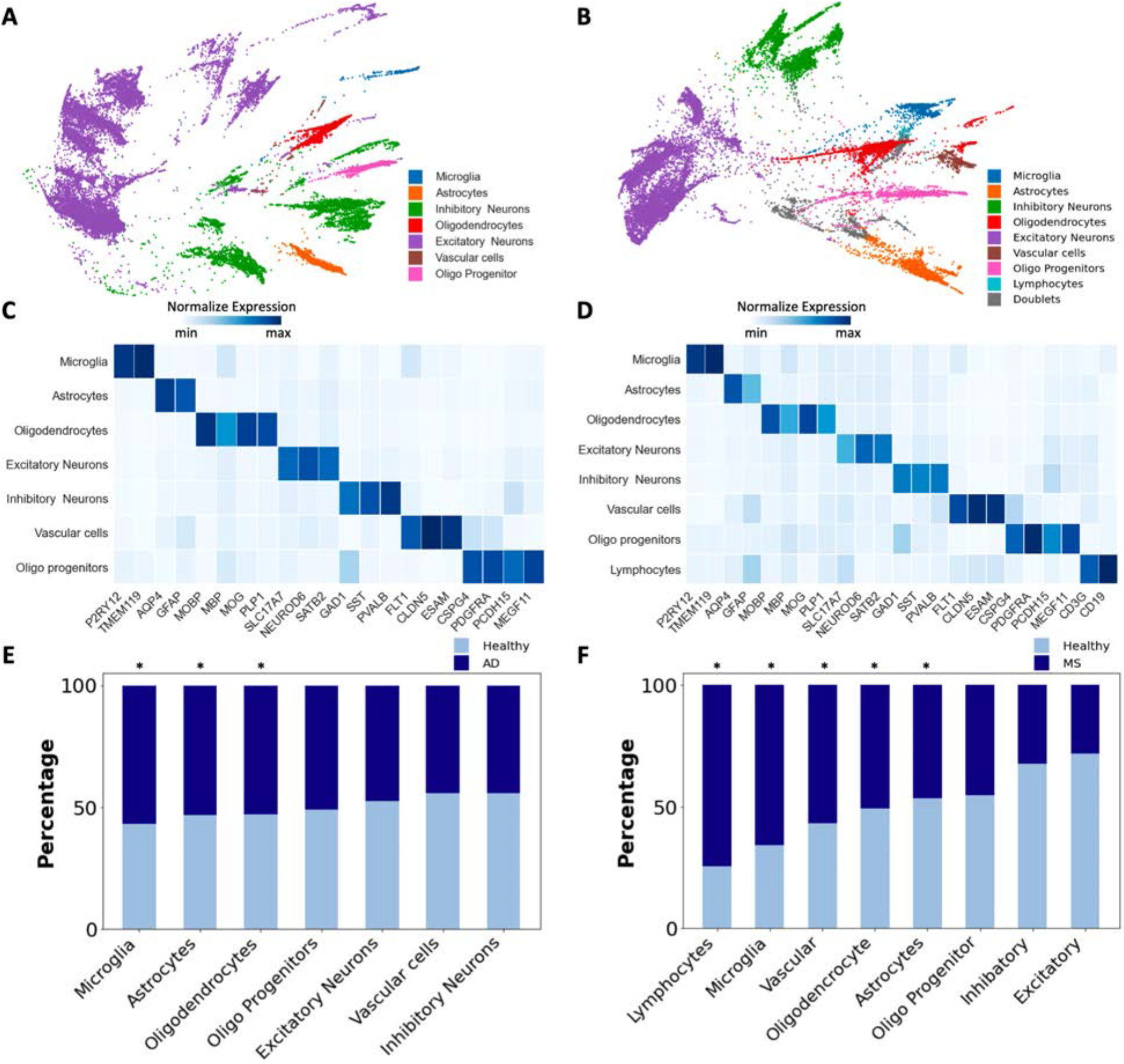
Diffusion condensation analysis of AD and MS snRNAseq data reveals enrichment and activation of microglia and astrocytes in disease. **(A)** 43,650 cells pooled from 48 AD patients and healthy donors. Samples were taken from disease free brain tissue and diseased brain tissue at early and late pathological stages. All major cell types were identified by diffusion condensation via persistence analysis of computed diffusion homology and visualized with PHATE [*18*]. (B) 46,796 cells pooled from 21 progressive MS patients and healthy donors. Samples were taken from disease free brain tissue and diseased brain tissue at acute and chronic stages of inflammation. All major cell types were identified by diffusion condensation via persistence analysis of computed diffusion homology and visualized with PHATE [*18*]. **(C)** Diffusion condensation reliably identifies cell types in AD brain tissue, as shown by average normalized expression of known cell type-specific marker genes. **(D)** Diffusion condensation reliably identifies cell types in MS brain tissue, as shown by average normalized expression of known cell type-specific marker genes. **(E)** Microglia and astrocytes are the most enriched cell types in AD using cross condition abundance analysis. **(F)** Microglia and astrocytes are significantly enriched in progressive MS using cross condition abundance analysis. E,F: p < 0.01, single-sided binomial test.

**Supplementary Figure 6:**
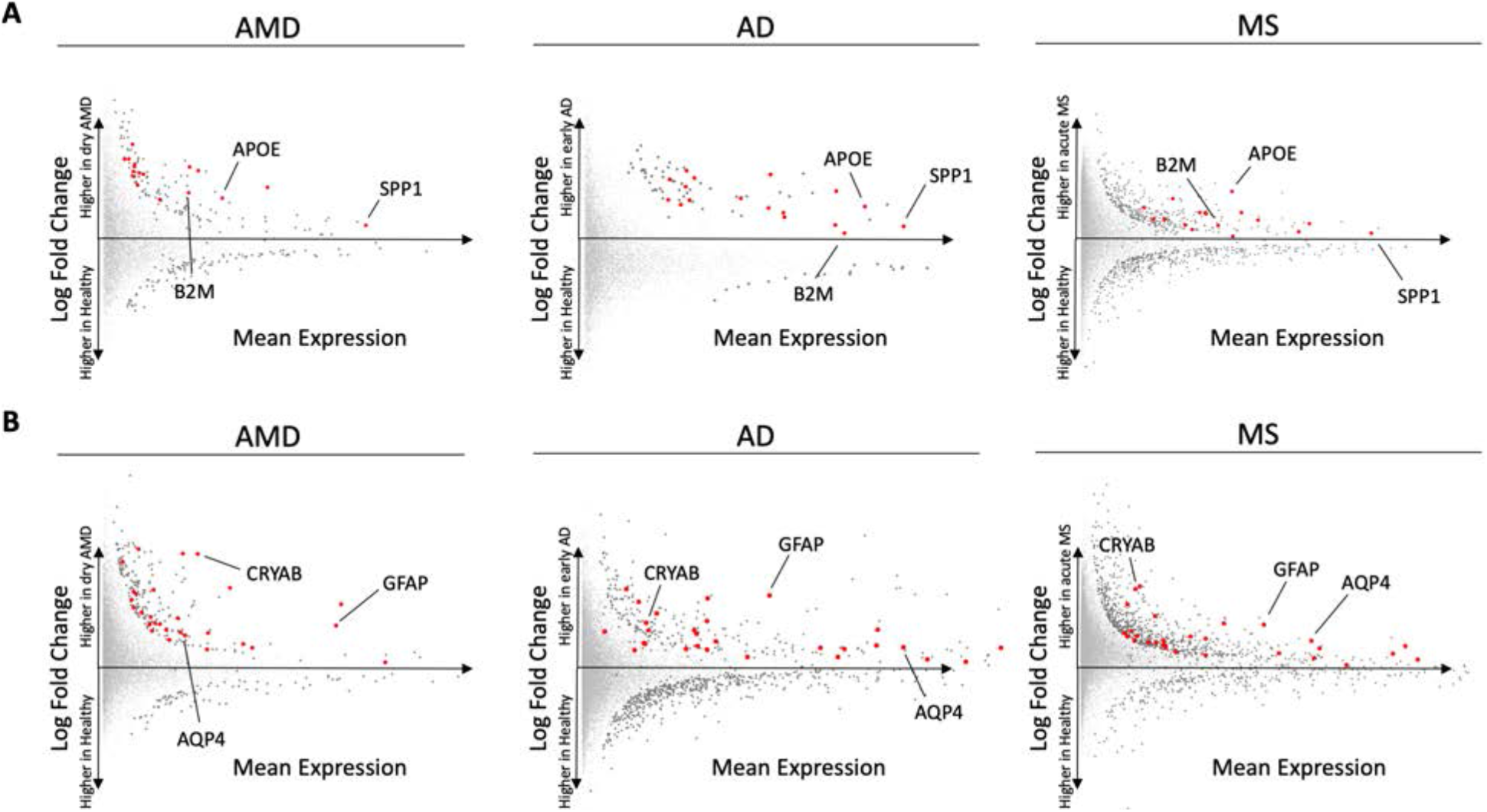
Cell type level differential expression analysis at coarse granularity across neurodegenerative diseases. **(A)** Performing differential expression analysis between microglia which originate from dry AMD patients or healthy controls identified a gene signature enriched in the early stage of dry AMD. By performing similar differential expression analysis between microglia from brain samples from patients with early AD, acute active MS, and controls, we identified a shared activation signature of 17 genes. This common signature includes *APOE*, *SPP1* and *B2M* while all other genes are highlighted in red. **(B)** Performing differential expression analysis between astrocytes which originate from dry AMD patients or healthy controls identified a gene signature enriched in the early stage of dry AMD. By performing similar differential expression analysis between astrocytes from healthy and early AD samples and healthy and acute inflammation MS samples, we identify a shared activation signature of 28 genes. This common signature includes *GFAP*, *AQP4*, and *CRYAB* while all other genes are highlighted in red.

**Supplementary Figure 7:**
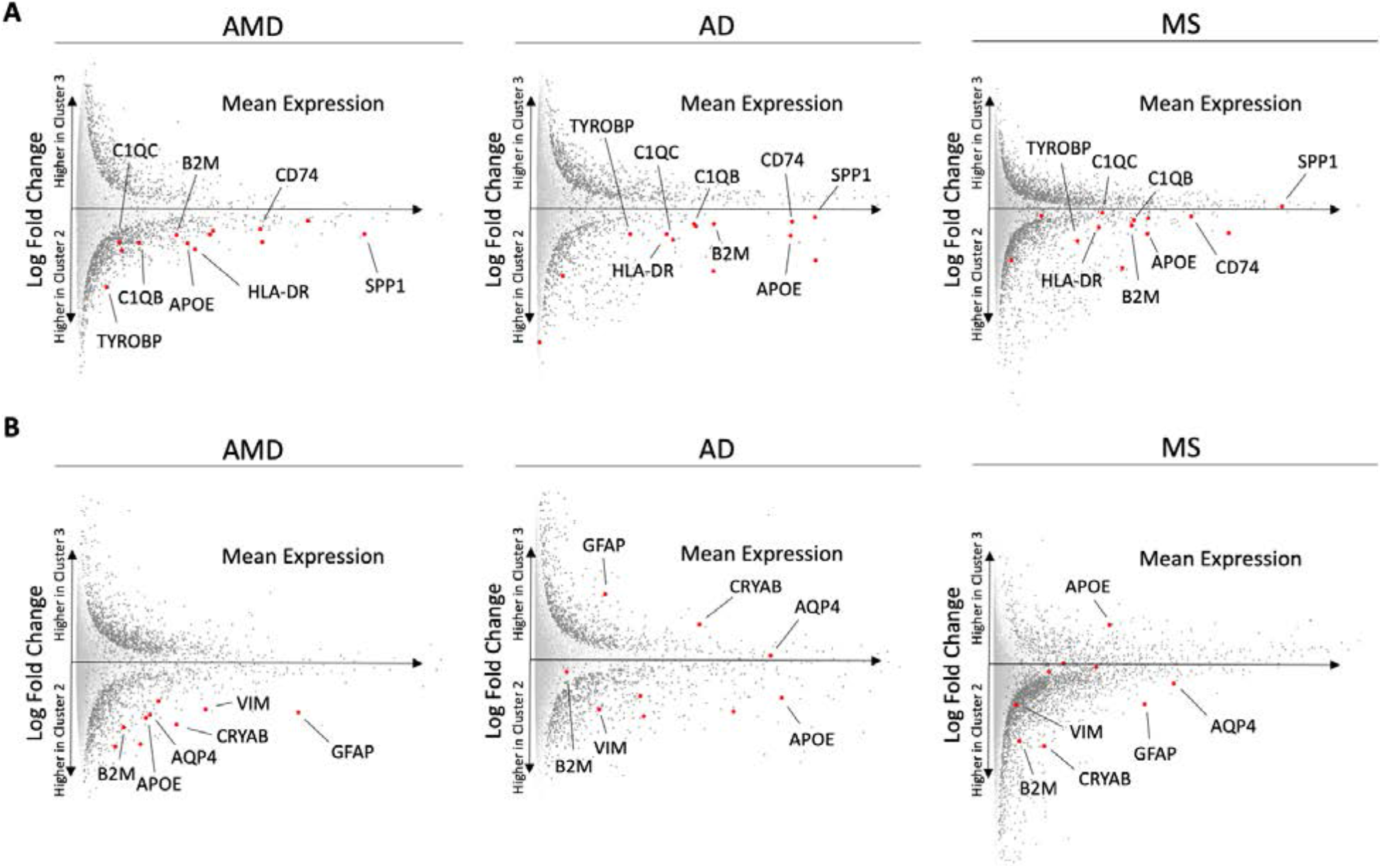
Shared microglial and astrocyte activation signature is diminished in advanced disease. **(A)** Comparing microglial cluster 3 (advanced or chronic inactive disease-enriched) to cluster 2 (early or acute active disease-enriched) revealed a significant reduction in the microglial activation signature across later stages neurodegeneration. **(B)** Comparing astrocyte cluster 3 (advanced or chronic inactive disease-enriched) to cluster 2 (early or acute active disease-enriched) revealed a significant reduction in the astrocyte activation signature across later stages neurodegeneration.

**Supplementary Figure 8:**
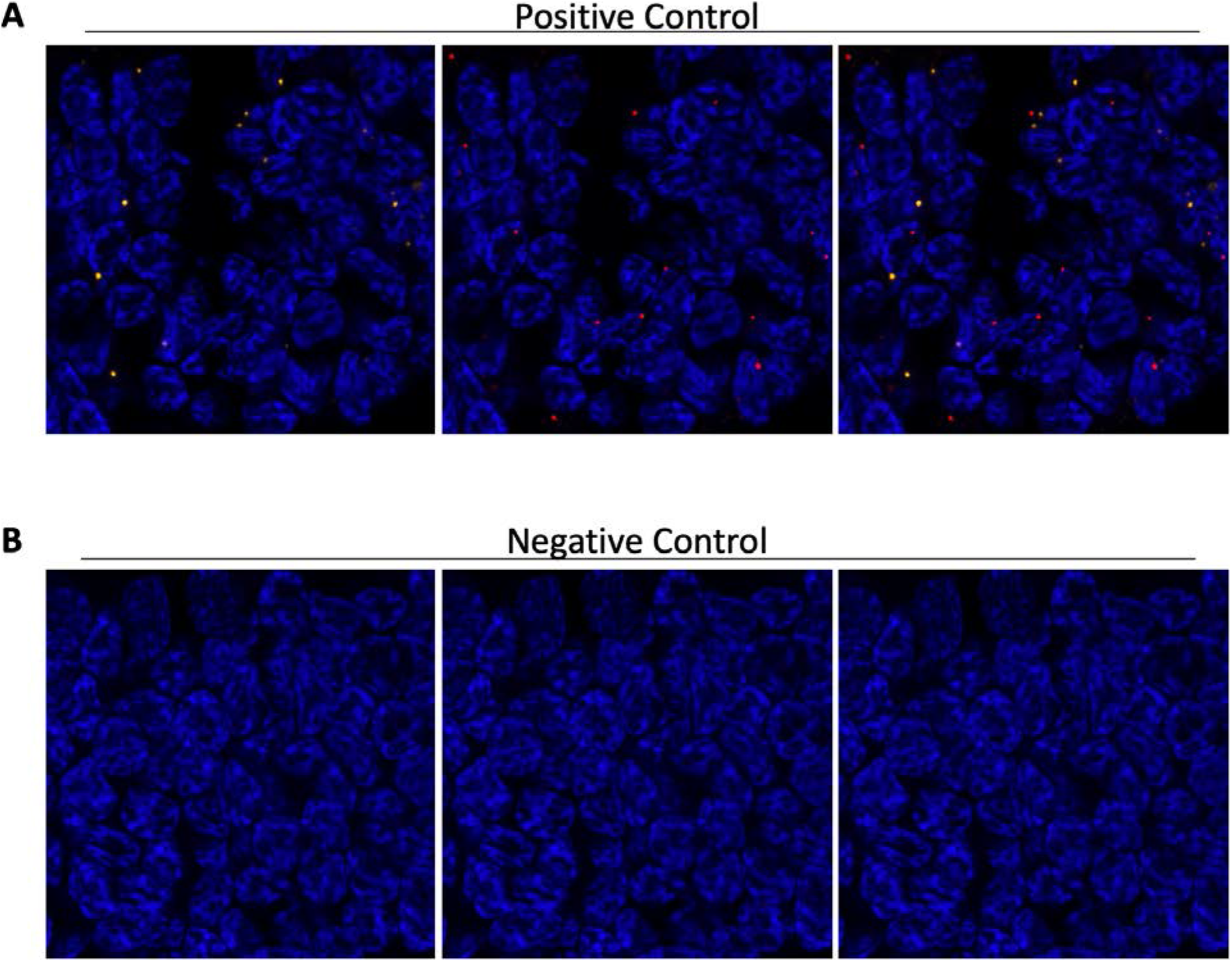
Positive and negative control probes for fluorescence in situ hybridization. Representative images of fluorescence in situ hybridization for **(A)** positive control probe (POL2RA labeled in red and UBC labeled in yellow) and **(B)** negative control probe (DapB labeled in yellow and red).

**Supplementary Figure 9:**
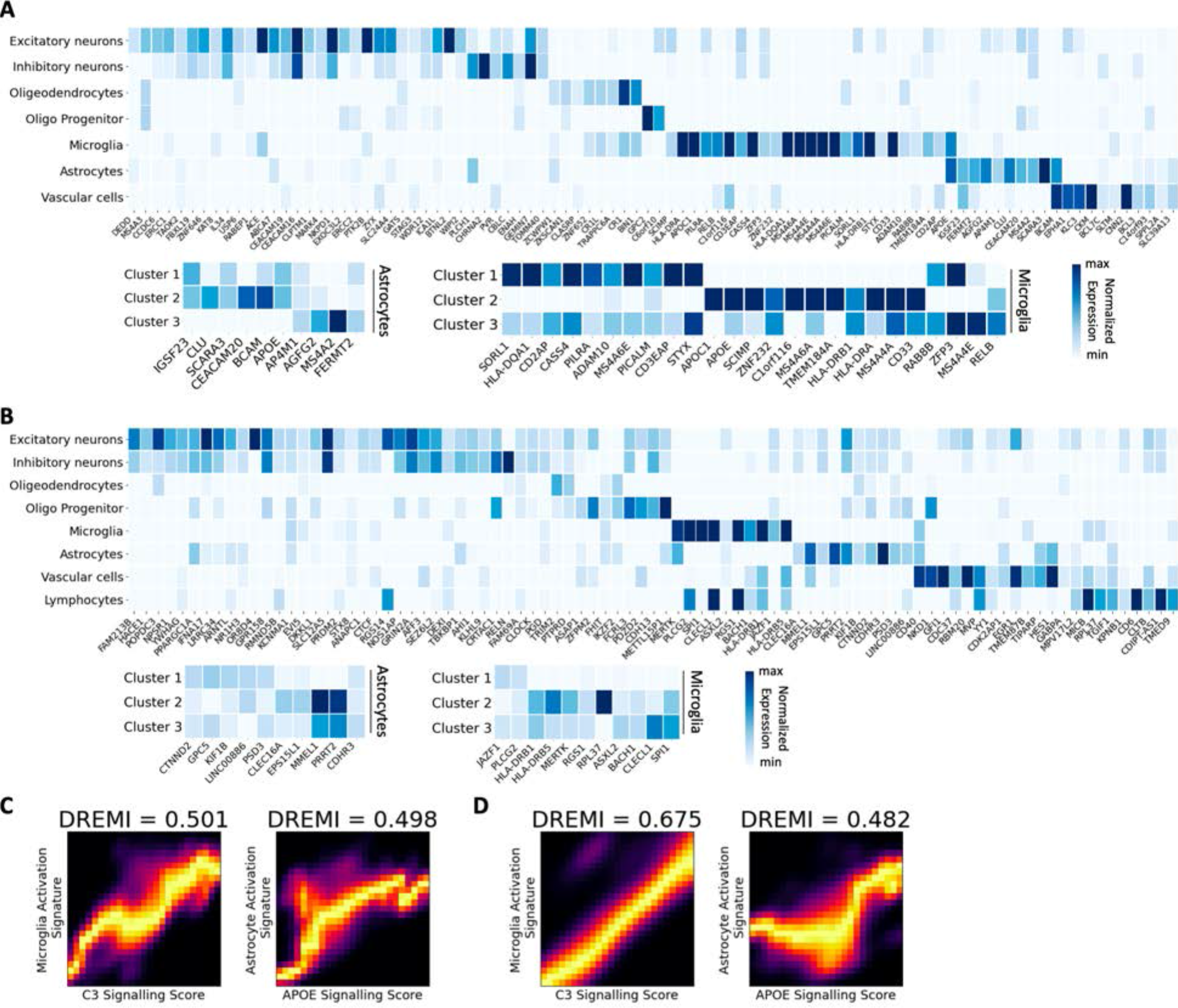
Mapping disease associated genes to cell types and cell states identified by diffusion condensation across granularities in AD and MS data. **(A)** Mapping AD risk associated genes [*44*] to diffusion condensation identified cellular populations across granularities for astrocytes and microglia. **(B)** Mapping MS risk associated genes [*45*] to diffusion condensation identified cellular populations across granularities for astrocytes and microglia. **(C)** DREVI plots of C3 and APOE signaling scores versus the composite activation signatures for microglia (left) and astrocytes (right) in AD. **(D)** DREVI plots of C3 and APOE signaling scores versus the composite activation signatures for microglia (left) and astrocytes (right) in MS.

**Supplementary Figure 10:**
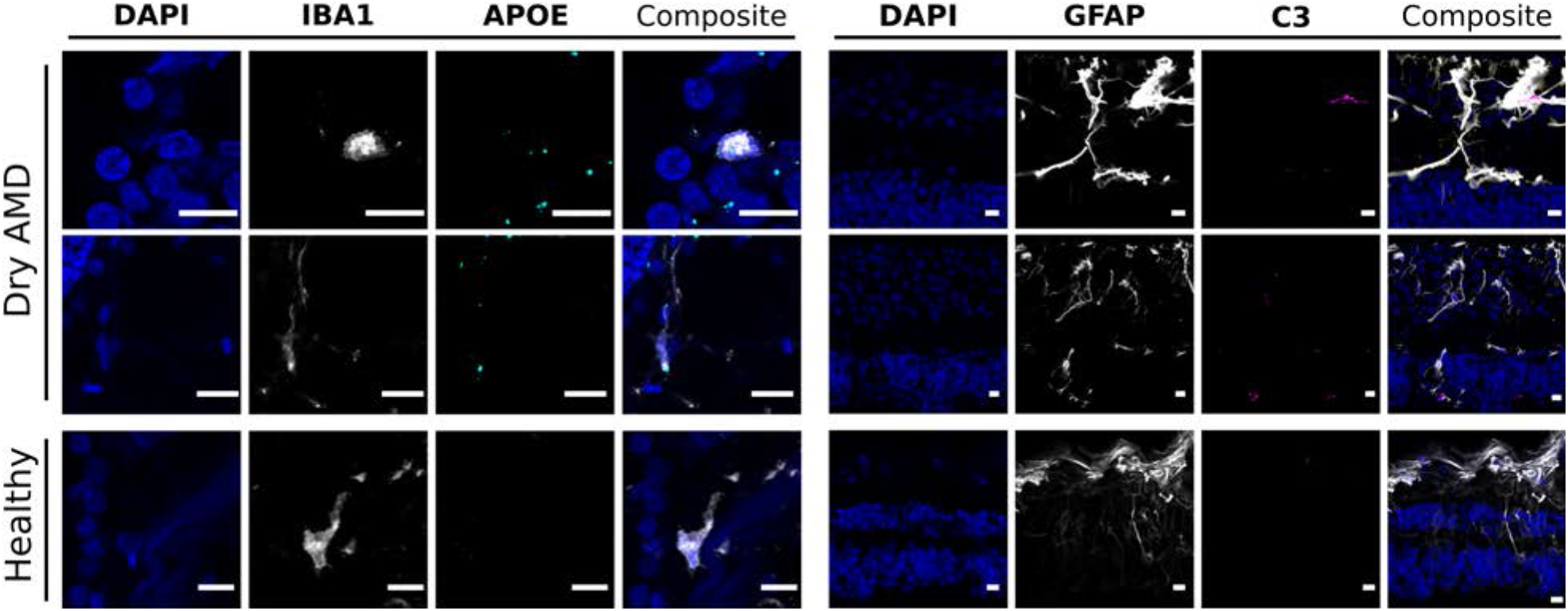
Microglia and astrocytes express elements of a recurrent signaling pathway. Representative images of in situ RNA hybridization of APOE (left; labeled in turquoise), C3 (right; labeled in pink) with immunofluorescence of microglial marker IBA1 or astrocyte marker GFAP (identified in white). Elevated expression of *APOE* is seen in IBA1-positive cells, and more abundant *C3* expression is seen in astrocyte-rich retinal layers from the from retina with dry AMD (top row) compared to healthy control (bottom row). All scale bars = 10μm.

